# Basal association of a transcription factor favors early gene expression

**DOI:** 10.1101/2024.03.26.586726

**Authors:** Sandrine Pinheiro, Mariona Nadal-Ribelles, Carme Solé, Vincent Vincenzetti, Yves Dusserre, Francesc Posas, Serge Pelet

## Abstract

Responses to extracellular signals via Mitogen-Activated Protein Kinase (MAPK) pathways control complex transcriptional programs where hundreds of genes are induced at a desired level with a specific timing. Gene expression regulation is largely encoded in the promoter of the gene, which harbors numerous transcription factor binding sites. In the mating MAPK pathway of *Saccharomyces cerevisiae*, one major transcription factor, Ste12, controls the chronology of gene expression necessary for the fusion of two haploid cells. Because endogenous promoters encode a large diversity of Ste12 binding sites (PRE), we engineered synthetic promoters to decipher the rules that dictate mating gene induction. Conformations of PRE dimers that allow efficient gene expression were identified. The strength of binding of Ste12 to the PRE and the distance of the binding sites to the core promoter modulate the level of induction. The speed of activation is ensured by favoring a basal association of Ste12 by using a strong dimer of PRE located in a nucleosome depleted region.

**Author Summary:** During development, cell fate decisions allow pluripotent cells to differentiate into various cell types. This process requires cells to integrate signals from their surroundings to initiate a complex transcriptional program. Budding yeasts can also undergo cell fate decisions. In presence of mating pheromones, haploid yeasts can activate a signaling pathway which can ultimately lead to the fusion of two haploid cells to form a diploid.

One transcription factor, Ste12, controls this mating transcriptional program. The promoters of these 200 upregulated genes display a large diversity in the organization of Ste12 binding sites. Therefore, it is challenging to decipher how Ste12 regulates the level and the timing of gene expression. To simplify this problem, we have generated synthetic promoters, where the configuration of Ste12 binding sites on the DNA can be controlled. We have identified which conformations of binding site dimers allow a functional association of the transcription factor. In addition, we have also shown that the basal association of Ste12 to the promoter is important for the fast gene induction. An unfavorable configuration of Ste12 binding sites or the presence of nucleosomes restrict the access of the transcription factor to the DNA and results in a slower expression.

## Introduction

*De novo* protein synthesis plays a central role in all cellular functions. This process can be controlled by internal regulatory inputs emanating from the cell cycle machinery(1,2), from fluctuations in circadian rhythms(3) or from oscillations in the metabolic state(4). In addition, extracellular cues such as stresses, nutrients or hormones can stimulate gene expression. In all eukaryotic cells, Mitogen-Activated Protein Kinase (MAPK) pathways play an essential role in transducing extracellular information into a cellular response, which generally includes the production of new proteins(5,6). The gene induction process may be transient, in order to adapt to a new stressful environment(7). However, if the protein production is sustained, it can profoundly modify the cellular physiology by altering its entire proteome(8). Cell fate decision systems implicated in cellular differentiation mechanisms rely on this *de novo* protein production to transform naive pluripotent cells into differentiated cells that will ultimately give rise to the different parts of a multicellular organism.

Despite their relative simplicity, unicellular organisms can also take complex decisions. The budding yeast *Saccharomyces cerevisiae* induces diverse transcriptional programs via MAPK cascades that allow this microorganism to make appropriate decisions in response to a specific stimulus. As an illustration, in low nutrient conditions, both haploid and diploid cells can alter their growth pattern to form pseudohyphae(9). Under more drastic nutrient limitations, diploid cells begin to sporulate(10). In rich medium and in the presence of a mating partner, haploid cells can commit to mating to produce diploid cells(11,12).

During the mating process, cells of opposing mating types (MATa or MAT α) communicate by secreting pheromones (a- or α-factor, respectively). At the cell surface, binding of pheromones stimulates a G-protein coupled receptor which in turn activates the three-tiered kinase cascade(11,12) (Supplementary Figure 1). The MAPKs, Fus3 and Kss1, release the inhibition of Dig1 and Dig2 on the transcription factor (TF) Ste12(13). This step is critical for activating the mating transcriptional program, resulting in the up-regulation of more than 200 genes(14). Note that a small fraction of these genes, which are typically cell-type specific, depend on co-activators such as Mcm1, a1 or α1 and α2 (15,16). The mating transcriptional response promotes the arrest of the cell cycle and the formation of the mating projection, two processes which are necessary to ensure a robust mating of the partner cells.

The proteins involved in the various stages of the mating process are tightly regulated in their expression levels and dynamics by Ste12. Genes involved in the early phase of mating, such as the establishment of a pheromone gradient (*BAR1*), MAPK signal transduction (*FUS3*, *STE12*), cell cycle arrest (*FAR1*) and cell agglutination (*AGA1*) are expressed rapidly after the detection of the pheromone(17). In contrast, genes involved in later stages, such as karyogamy (*KAR3*) and membrane fusion (*FIG1*) will be expressed with a delay(17,18). The mechanisms which allow a single transcription factor (Ste12) to orchestrate this chronology of gene expression remain poorly understood.

The promoter sequence upstream of the protein-coding sequence regulates the level and dynamics of transcription. The promoter combines two distinct segments: the core promoter and the regulatory region(19,20). With a typical length of 100-200bp, the core promoter contains the TATA box which is recognized by the TATA-binding protein. This protein recruits other general transcription factors and contributes to the formation of the Pre-Initiation Complex (PIC). In yeast, the regulatory region is typically smaller than 1kb and carries Upstream Activation Sequences (UAS) recognized by TFs. Transcriptional activation is induced via the Mediator complex, which bridges the TFs and the RNA polymerase II, thereby assembling the PIC to initiate transcription(21,22).

Studies have shown that induction of mating genes requires the formation of a Ste12 homodimer on the UAS of mating promoters(23,24). The region is recognized by the protein via the minimal DNA consensus motif TGAAAC, commonly referred to as the Pheromone Response Element (PRE)(25–27). In addition to these consensus PREs, Ste12 also binds with lower affinity to non-consensus sites, known as PRE-like sites. In general, multiple PREs or/and PRE-like sites can be identified on the promoters of mating genes(17,28) and the arrangement of these PRE motifs controls the expression profile of a gene(17,27). Unfortunately, defining simple rules that would allow to predict the expression pattern of mating genes is challenging because of the wide diversity in configurations present on endogenous promoters. In addition, the identification of all possible Ste12 binding sites is difficult, since it is unclear how much a PRE-like site can deviate from the consensus while retaining affinity for Ste12.

Obviously, a complex promoter architecture is not restricted to mating genes. Essentially, all endogenous promoters harbor an intricate arrangement of TF binding sites, often combining multiple TF inputs(29). General rules determining gene expression patterns have been obtained by analyzing libraries of synthetic promoters tested for a wide diversity of binding site organization(30,31). In general, the number of binding sites, their affinity and their distance from the core promoter can all influence the expression output. A prolonged residence time of a TF on a promoter will increase the expression output, as the chance of initiating transcription via the recruitment of the Mediator complex and formation of the PIC will rise(32). However, it is not known whether a high transcriptional output is necessarily correlated with fast gene expression of whether these two parameters (i.e. strength and speed of induction) can be decoupled.

To decipher how the promoter sequence modulates gene expression dynamics, we have engineered synthetic promoters under the control of the TF Ste12. These promoters combine different PRE conformations to understand the parameters that regulate the transcriptional program for cell fate decisions during mating. The dynamics and level of protein production were quantified to assess how the orientation and the spacing between PRE pairs, their affinity and their location along the promoter influence the transcriptional response. Mutating the PRE consensus to lower the affinity of Ste12 for the promoter or changing the location of the sites on the promoter influenced predominantly the level of induction. However, our data show that the ability of Ste12 to associate to a promoter prior to the stimulus either by controlling by the Ste12 binding site conformation or the access to the DNA in nucleosome depleted regions favors faster gene induction.

## Results

To measure the dynamics of gene expression in the mating pathway, we use a dynamic Protein Synthesis Translocation Reporter (dPSTR)(33). It consists of two transcriptional units: the first one encodes a fluorescent protein fused to a synthetic “zipper” coiled-coil helix (SynZip)(34) and is expressed constitutively. The second unit encodes two NLSs linked to a complementary SynZip and is placed under the control of a promoter of interest. Because SynZips form strong heterodimers, upon the expression of the NLS, the fluorescent protein relocates into the nucleus (Figure 1A and B and Supplementary Figure 2A and B). This sensing strategy allows a rapid quantification of protein production, which can otherwise be impaired by the slow maturation kinetics of fluorescent proteins.

**Figure 1.**
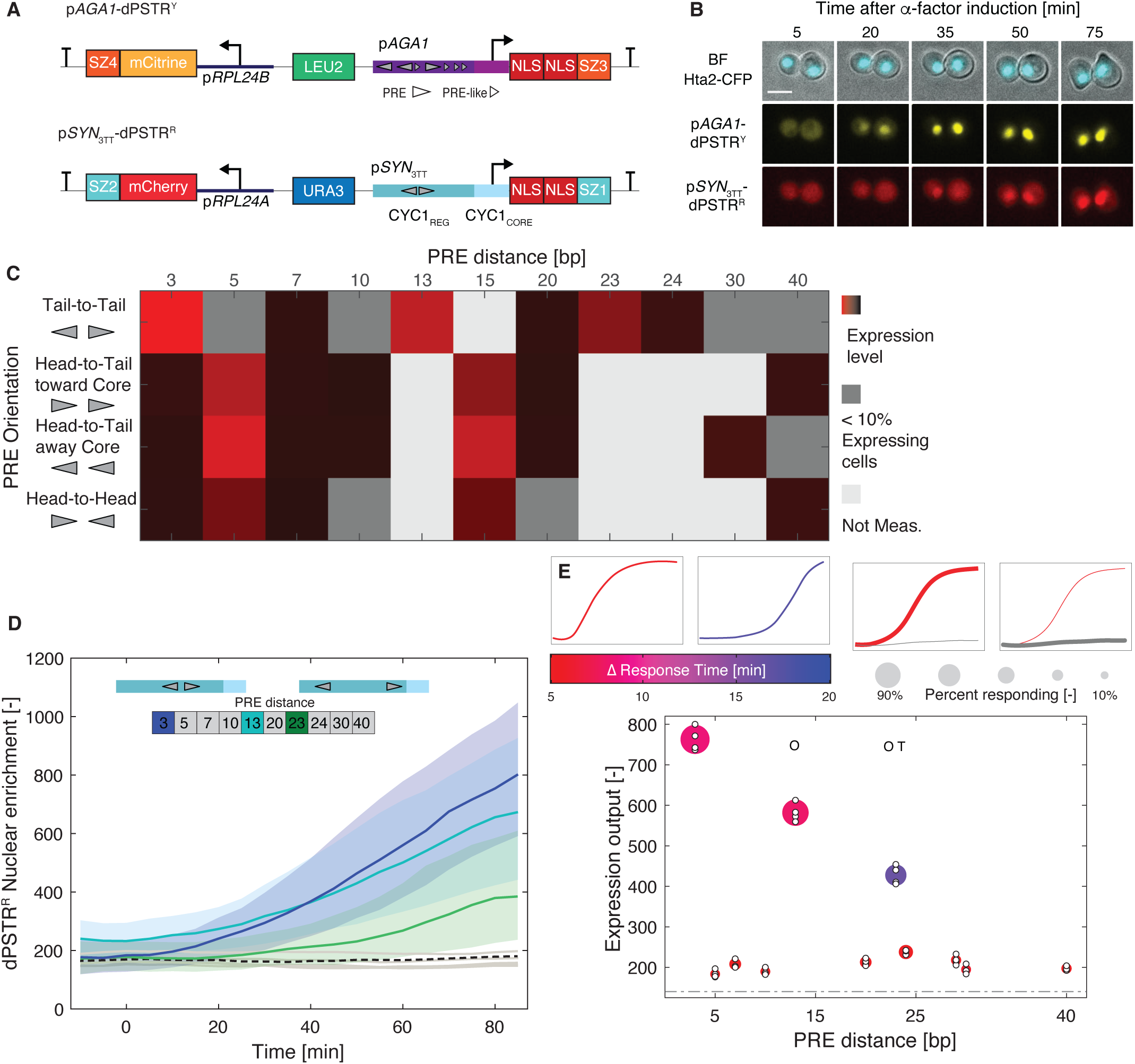
Effect of the orientation and spacing of Ste12 binding sites on the expression output. A. Scheme of the two reporters present in the strains. The reference p*AGA1*-dPSTR^Y^ and a test construct, here the p*SYN*3TT-dPSTR^R^. Both constructs encode the production of a constitutively expressed fluorescent protein and an inducible peptide containing two nuclear localization sequences (NLS) and a SynZip (SZ1 or SZ3, which can form a strong heterodimer with SZ2 and SZ4, respectively). The reference dPSTR^Y^ is under the control of the *AGA1* promoter in the yellow channel and the test dPSTR^R^ is regulated by a synthetic promoter of interest based on a modified *CYC1* promoter. B. Images of cells induced with 1µM α-factor at time 0. The nuclear enrichment of the fluorescent proteins serves as a measure of promoter activity. The scale bar represents 5 µm. C. Matrix representing the mean expression output for p*CYC1* containing two PRE sites with four different orientations and spaced between 3 to 40 bp. The strength of the red color is proportional to the expression output of the promoter. Dark gray areas represent construct where fewer than 10% of cells overcome the expression threshold. Light gray squares are PRE conformations that were not measured. D. Time course of the nuclear enrichment of the dPSTR^R^ for various distances of PRE placed in tail-to-tail orientation. The three functional conformations are plotted in blue (3bp) light blue (13bp) and green (23bp). The solid lines represent the median and the shaded area, the 25- to 75- percentile of the population. Gray lines represent the median of non-functional PRE conformations. The black dashed line is the median of the control synthetic promoter without PREs inserted. E. Summary graph displaying the expression output, the speed and the fraction of responding cells for various spacings of the PRE dimer placed in the tail-to-tail orientation. The color of the marker indicates the difference in response time between the synthetic promoter and the reference p*AGA1*-dPSTR^Y^, with fast responding promoters in red and slow ones in blue as indicated by the two small schematic graphs on the left. The size of the marker represents the fraction of responding cells as depicted by the two schemes on the right. The expression output of individual replicates is indicated by small white dots. The expression threshold based on the level of p*SYN*3TT is indicated by the dashed dotted line. The letters O and T indicate a significant difference between the mean of the replicates (t-test: p-val < 0.05) in the timing of induction (T) or in the expression output (O) relative to the p*SYN*3TT.

In our assays we combine two dPSTRs. In the yellow channel, we have a reference p*AGA1*- dPSTR^Y^. *AGA1* encodes an agglutinin that promotes cellular adhesion of mating pairs. This gene has been shown to belong to the early gene category and is expressed at high levels, similarly to the well-established p*FUS1* reporter(17,26,35,36). In the red channel, we monitor the dPSTR^R^ controlled by a promoter of interest (Figure 1A and B). The expression outputs of a dPSTR^Y^ and a dPSTR^R^ controlled by the same p*AGA1* promoter are tightly correlated (Supplementary Figure 2C). While p*AGA1* is activated in the entire population, other promoters are activated in a fraction of the population (Supplementary Figure 2D, E and F). Importantly, the comparison of the nuclear enrichment dynamics between the reference dPSTR^Y^ and the test dPSTR^R^ provides an accurate measurement of the kinetics of activation of our promoter in each cell. Notably, it overcomes cell to cell fluctuations due to the cell-cycle regulated activation of the mating (Supplementary Figure 2G and H).

### Construction of a pheromone−inducible synthetic promoter

Starting from the *AGA1* promoter, we first selected an alternative core promoter based on the *CYC1* promoter (Supplementary Figure 3). Although the *CYC1* core ensures a sizable inducibility, the expression dynamics are delayed by 5 minutes relative to p*AGA1*. This delay reinforces the idea that the core promoter identity contributes to define both the level and kinetics of gene expression(17). In all our synthetic promoters, we will use the same p*CYC1* core promoter allowing to focus our study on the role of the regulatory sequence in the dynamics of gene expression.

The promoter activity is controlled by the regulatory region, which can contain multiple TF binding sites. In p*AGA1*, we have identified three consensus PREs and multiple PRE-like (Supplementary Figure 3A). We had previously determined that two PREs spaced by 29 bp were important for the fast and high induction of the promoter upon α-factor treatment(17). To test if these two PRE sites spaced by 29 bp are sufficient to control gene induction, we placed them in a completely synthetic context. To do so, we have selected a *CYC1* promoter, which was modified to decrease nucleosome binding(37) and where identified TF binding sites were mutated(29), as well as sequences that resembled potential PRE sites. Two PREs spaced by 29 bp placed in this synthetic context are not sufficient to induce expression (Supplementary Figure 3B and C). The inducibility of the construct is recovered when the original sequence between the two PREs from p*AGA1* is included. This difference was attributed to the presence of a PRE-like site located 3bp away from the second consensus PRE(24). Engineering a *pCYC1* UAS containing two PRE-consensus sites 3bp apart fused to the *pCYC1* core, resulted in a rapid pheromone-inducible synthetic promoter (Supplementary Figure 3B, C and D). This initial synthetic construct (p*SYN*3TT- 2PRE spaced by <u>3</u> bp in <u>T</u>ail-to-<u>T</u>ail configuration) will be used as a reference for the systematic alterations that will be performed on the PRE sites to generate our library of Ste12-dependent promoters.

### PRE site conformations

Our initial efforts to engineer a synthetic promoter have clearly demonstrated that while two PREs are required to induce gene expression, not all PRE conformations result in a functional promoter. Endogenous promoters harbor a wide diversity of PRE configurations, and it is difficult to predict which ones of these PRE or PRE-like sites contribute to the overall expression output by allowing formation of a Ste12 dimer. Dorrity *et al.* have described the canonical 3bp tail-to-tail conformation (as found in our p*SYN*3TT) as the most favorable binding *in vitro* for the Ste12 DNA-binding domain (DBD) fragment(24). This PRE conformation is found on multiple endogenous promoters such as p*AGA1*, p*STE12,* p*KAR4* and p*FAR1*. However, many endogenous promoters don’t harbor this element. Therefore, it is likely that other conformations can promote a functional Ste12 binding. In addition, Su *et al.* demonstrated that a head-to-tail conformation could induce gene expression, while a head-to-head positioning prevented Ste12 binding when placed in close proximity(27).

To identify functional PRE conformations, we have systematically altered the arrangement of two PRE sites in our synthetic promoter construct (Figure 1C). With PRE in tail-to-tail orientation, the promoter becomes non-functional if the distance between the PREs is extended to 5, 7 or 10 bp. Interestingly, transcriptional activity is recovered when the spacing is set at 13 bp. This distance can even be further extended to 23 bp, resulting in a low and slow expression output (Figure 1D and E). These functional binding conformations can be rationalized by the fact that the DNA-helix turn corresponds to 10.5 bp(38). Therefore, spacing the two PRE by 3, 13 or 23 bp, positions the two Ste12 proteins in a similar interaction geometry. The 13bp tail-to-tail conformation is found in the promoter of *SST2*, however, the PRE spacings of 23 was not identified in the endogenous promoters we inspected.

If the orientation of the PRE is changed to head-to-tail or head-to-head conformations, the preferred distance between PREs becomes 5 bp (Figure 1C, Supplementary Figure 4). A spacing of 15 bp is also transcriptionally active, but other spacings result in no or very low expression output. The head-to-tail conformation spaced by 5 bp is readily found in numerous promoters (p*AGA1*, p*FUS1*, p*PRM1*, p*FIG1*), while the 15bp seems less prevalent (p*FUS2*). In contrast, the 5 bp head-to-head conformation does not seem to be frequent because we did not identify it in the two dozen endogenous promoters investigated. We also verified by deleting *STE12* that the induction of these synthetic promoters was strictly dependent on this transcription factor (Supplementary Figure 3E and F).

Overall, these findings validate the use of synthetic promoters to identify functional binding site conformations. These results can be transferred to endogenous promoters to pinpoint the sites implicated in the mating-dependent induction. The promoter sequences can be analyzed to identify short-range interaction with specific spacings of 3bp for tail-to-tail orientations and 5 bp for the other orientations while including a possible increment of 10 or 20 bp corresponding to one or two DNA helix turns.

### Contribution of Ste12 activation domain to gene expression

Ste12 activates transcription via specific PRE conformations. This extended set of functional PRE pairs indicates a surprising flexibility from Ste12 to homodimerize (Figure 2A). *In vitro* data have shown that the DBD of Ste12 can dimerize to bind to two PREs(24). However, it is difficult to imagine how the Ste12 DBD can support the various dimerization conformations identified with our synthetic promoters, which include diverse orientations and distances. Since it has also been shown that the activation domain (AD) of Ste12 can multimerize(23), we wanted to determine if this flexible region of the protein contributes to the stabilization of some of the Ste12 binding conformations that we have identified, for instance, on PREs separated by larger distances.

**Figure 2.**
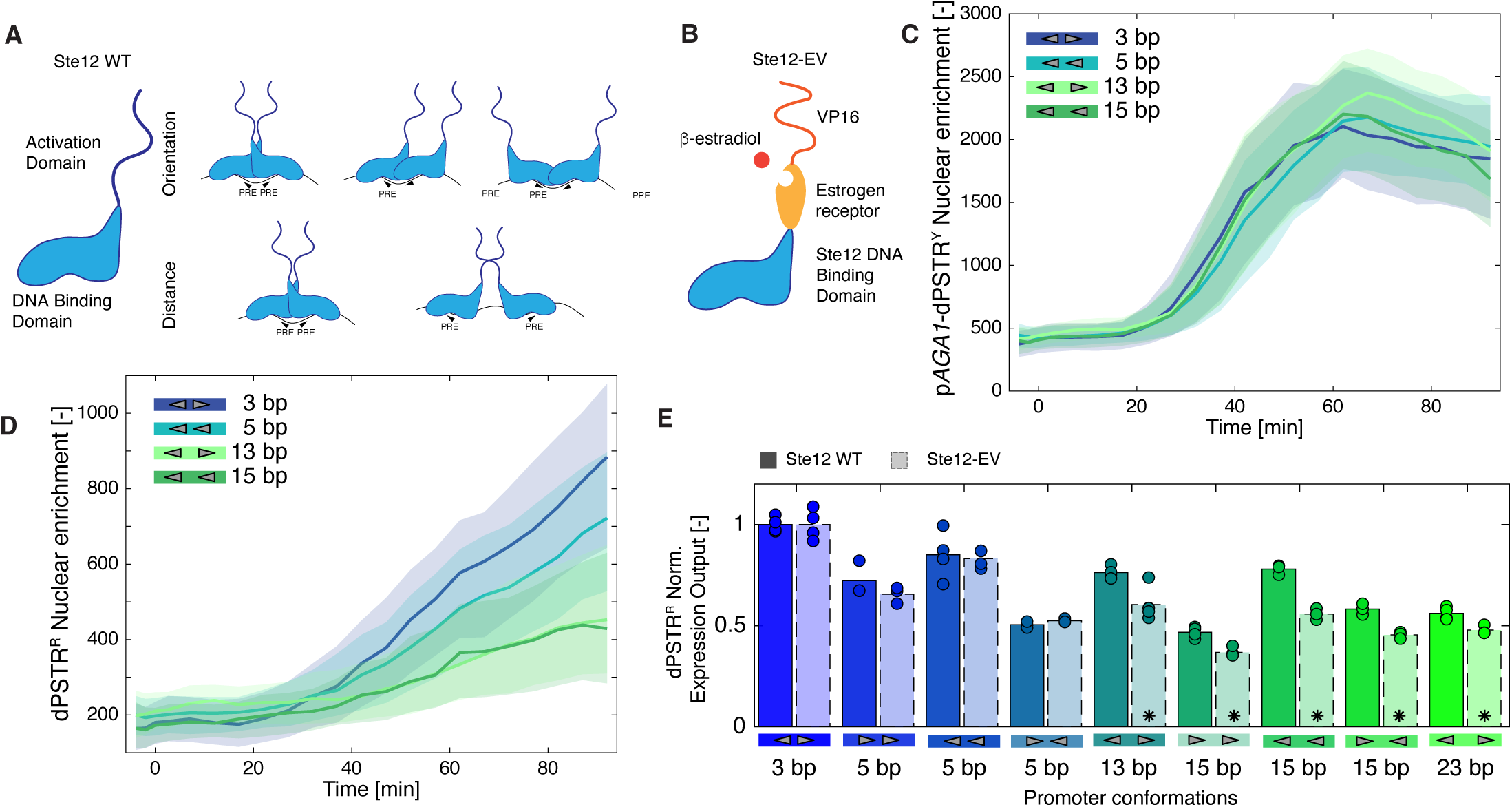
Dominant role of Ste12 DNA binding domain for PRE conformation selection. A. Schematic of the Ste12 transcription factor composed of a DNA-binding domain and a flexible activation domain. The small schemes describe the diversity of Ste12 homodimer interactions that could take place when varying the orientation or the distances of the PRE sites on the promoter. B. Scheme of the Ste12-EV chimeric transcription factor where the activation domain of Ste12 has been replaced by the estrogen receptor and the VP16 activation domain resulting in an β-estradiol responsive TF. C., D. Dynamics of nuclear relocation following stimulation with 1µM β-estradiol at time 0 for strains containing the p*AGA1*-dPSTR^Y^ (C) and various p*SYN*-dPSTR^R^ (D). The solid lines represent the median of the population and the shaded area, the 25- to 75- percentiles. E. Comparison of the inducibility of various PRE conformation with the Ste12 WT or the Ste12- EV (black borders). The Expression Outputs (EO) for the synthetic reporters induced by the Ste12 WT or the Ste12-EV were normalized relative to the EO of the reference p*SYN*3TT. The bar represents the mean response of the replicates shown by the circles. A significant difference between the normalized EO of Ste12-WT and Ste12-EV based on the mean of the individual replicates is indicated by a star (t-test: p-val < 0.05)

To test the relative contributions of the DBD and the AD of Ste12 for the inducibility of our various synthetic promoters, we replaced the region coding for the activation domain in the endogenous *STE12* locus by a fusion between the human estrogen receptor and the VP16 activation domain (EV)(39,40). Using this construct (Figure 2B, Ste12-EV), Ste12-responsive genes will be activated by stimulating the cells with β-estradiol, which promotes the relocation of the Ste12-EV to the nucleus and by-passes the mating MAPK cascade. The strength of induction of the PRE containing promoters will be governed only by the binding of the Ste12 DBD. Indeed, no interaction from the EV domain is expected (Supplementary Figure 5 A and B). Thus, we can test if the absence of the Ste12 activation domain lowers the inducibility of our synthetic promoters.

Stimulating the cells expressing Ste12-EV with β-estradiol results in a potent, while relatively slow, activation of the p*AGA1*-dPSTR^Y^ and of the p*SYN*3TT-dPSTR^R^ (Figure 2C and D). The time required to relocate Ste12 from the cytoplasm to the nucleus may contribute to this delay(40,41).

While the p*SYN*3TT-dPSTR^R^ is strongly induced in the Ste12-EV background, synthetic promoters bearing no PRE or a single PRE fail to be induced (Supplementary Figure 5 C and D). Similarly, if the PREs are spaced by 40 bp, no relocation of the dPSTR^R^ can be observed. These control experiments demonstrate that the Ste12-EV chimeric transcription factor relies solely on the DBD domain of Ste12 for activation and if the promoter contains a single PRE or 2 PREs placed in an undesired configuration, the association of Ste12-EV to the DNA is too weak to promote transcription, despite the presence of the potent VP16 activation domain.

To determine if the activation domain of Ste12 plays a specific role in the capacity of Ste12 to promote transcription for a subset of PRE conformations, we compared the strength of induction of various synthetic promoters from wild-type Ste12 and from Ste12-EV (Figure 2E). The expression output was normalized relative to the induction of the p*SYN*3TT-dPSTR^R^. Significantly lower induction level of the dPSTR^R^ was observed for the Ste12-EV when the two PREs are spaced by 13, 15 or 23 bp. Note that the normalized p*AGA1*-dPSTR^Y^ expression output and the fraction of responding cells remain similar between Ste12 and Ste12-EV (Supplementary Figure 5E and F). These results suggest that when the two PREs are spaced by more than 5 bp, the activation domain of the transcription factor plays a more important role in promoting the expression of the downstream gene, possibly by stabilizing the formation of the Ste12 dimer when the conformation of the two PRE is not optimal.

### Affinity of the PRE sites

Suboptimal PRE-arrangements can prevent Ste12 from binding to the promoter and limit the expression output. However, even in the context of PRE-dimers with optimal arrangements, Ste12 binding on mating promoters is affected if the binding site carries point mutations (PRE-like) as is found in a majority of endogenous promoters which harbor a combination of a consensus PRE and a PRE-like site (p*AGA1*, p*FIG1*)(24). A single base change to the TGAAAC consensus sequence can have a very different effect on the Ste12 affinity. While TaAAAC has only a 20% decrease in affinity, TGAgAC or TcAAAC result in a more than 95% reduction in competitive binding relative to the consensus sequence(27).

Therefore, to test the influence of the strength of the PRE site on the expression output, two PREs spaced by 3 base pairs were used and the sequence of one of the binding sites was modified. A clear decrease in the fraction of responding cells and in the level of expression are observed for the six PRE-like variants tested (Figure 3A and B). This is in line with previous measurements where the lowering of the affinity of the DNA-binding site results in a weaker gene expression output(27,30). We can imagine that the lower affinity of the site decreases the residence time of Ste12 on the promoter and therefore limits the expression output. Importantly, however, the dynamics of gene expression is not influenced by the lowering of the binding site affinity and all the promoters tested here, despite their lower induction level, are induced rapidly (Figure 3D).

**Figure 3.**
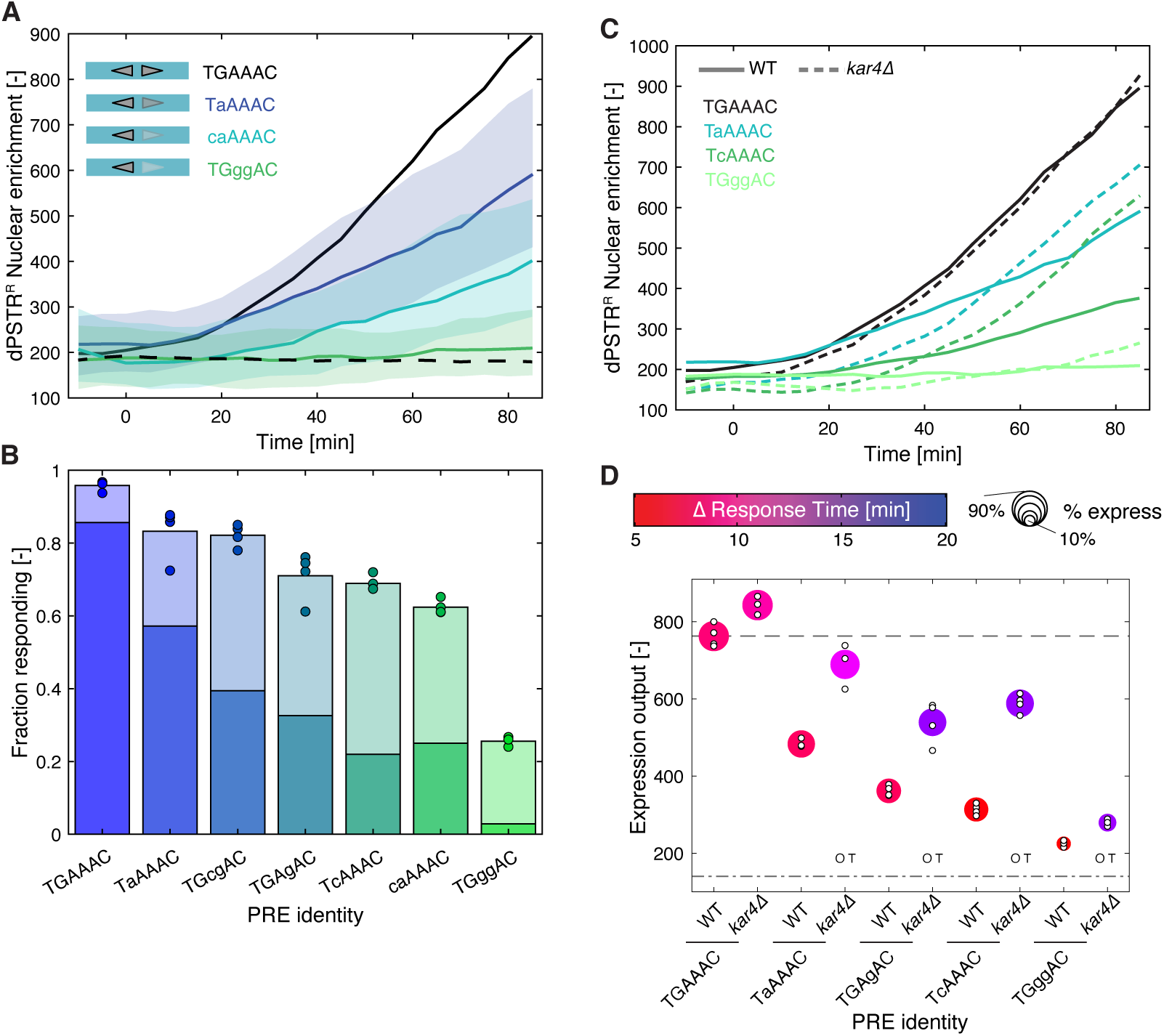
Influence of Kar4 deletion on low affinity PRE sites. A. Dynamics of nuclear relocation for the dPSTR^R^ controlled by two PRE spaced by 3 bp in tail- to-tail orientation, where one of the PRE sequences was mutated to alter its binding affinity (colored lines: median of the population and shaded area: 25- to 75- percentile). The solid black line represents the median response of the consensus PRE (p*SYN*3TT), while the dashed line represents the control promoter without PRE. B. Fraction of responding cells for the various PRE sequences. The darker portion of the bar represents the fraction of highly expressing cells (expression output > 50% of reference expression output) and the light bar the fraction of low-expressing cells (> 20% expression output < 50%). The marker represents the fraction of responding cells for individual replicates. C. Dynamics of nuclear enrichment of WT (solid lines) and *kar4Δ* cells (dashed lines) with a synthetic promoter with 2 PREs spaced by 3 bp in tail-to-tail orientation and where one of the PRE sequences has been mutated to decrease the affinity to Ste12. D. Summary graph displaying the expression output, the speed and the fraction of responding cells for two PRE spaced by 3 bp in tail-to-tail orientation and where one of the PRE sequences has been mutated to decrease the affinity to Ste12 in WT and *kar4Δ* cells. The color of the marker indicates the difference in response time between the synthetic promoter and the reference p*AGA1*- dPSTR^Y^. The size of the marker represents the fraction of responding cells. The expression output of individual replicates is indicated by small white dots. The dashed line represents the expression output and the dashed dotted line the expression threshold calculated based on the p*SYN*3TT. The O and T indicate a significant difference between the mean of the replicates (t-test: p-val < 0.05) in the timing of induction (T) or in the expression output (O) between the WT and *kar4Δ* strains for the same promoter.

As hypothesized in a previous study, the association of Ste12 on promoters containing suboptimal PRE-arrangements seems to be enhanced by Kar4(17). Indeed, late pheromone responsive promoters like p*FIG1* show minimal expression in *kar4Δ* cells, while fast ones are independent of Kar4. Therefore, to evaluate the contribution of Kar4 to the expression output of synthetic promoters containing PRE sites with different affinities for Ste12, we measured a set of promoters in *kar4Δ* cells. On promoters containing a PRE-like site, our measurements suggest that Kar4 has a dual role. On the one hand, Kar4 contributes to the rapid activation of Ste12-bound promoters, while, on the other hand, it limits the level of induction of the promoter (Figure 3C and D and Supplementary Figure 6A and B). A promoter with two PRE spaced by 13bp in tail-to-tail orientation follows the same trend of slow but high induction in *kar4Δ* cells, while the promoters with 5 and 15bp in head-to-tail configuration are minimally affected by the same deletion (Supplementary Figure 6C and D). Moreover, the high induction observed in *kar4Δ* cells for the promoter with a PRE-like seem to act via the activation domain of Ste12 because in the Ste12-EV chimeric protein, we don’t see an effect of the deletion of *KAR4* (Supplementary Figure 6E and F).

Taken together, these results indicate that the combination of a PRE with a PRE-like on a promoter tends to decrease the transcriptional output compared to two PRE sites, while the speed of gene induction remains unaffected, thanks to the contribution of Kar4. However, the phenotype associated with the deletion of KAR4 is complex because in the absence of this gene, the dynamics of induction of our synthetic promoters are slowed down, while their expression output is increased.

### Location of PRE sites on the promoter

In endogenous promoters, the distance of the PRE sites relative to the core promoter can vary greatly from -150 (p*FUS1*) to -440 p*STE12*. To test the influence of this parameter, we have moved the two PREs from p*SYN*3TT from their original position at -223 from the Start codon between -183 to -413 bp. Extending the distance between the Ste12 binding sites and the core promoter leads to a general decline of the expression output of the promoter (Figure 4 A and B). A similar behavior has been observed for the Msn2 stress response TF where moving its binding sites away from the core promoter lowered the expression output of the promoter(42). Moreover, we demonstrate that this lower expression output is not correlated with a decrease in the speed of gene expression, since all the p*SYN* tested remain fast (Figure 4C). Many of these promoters are in fact slightly faster than the reference p*SYN*3TT, possibly because they have an increased basal expression level. In these synthetic constructs, the binding dynamics of Ste12 should not be modified by the location of the PREs on the promoter. Thus, we speculate that the increased distance between the Ste12 binding sites and the core promoter precludes the ability of Ste12 from activating the general transcription factors via the Mediator.

**Figure 4.**
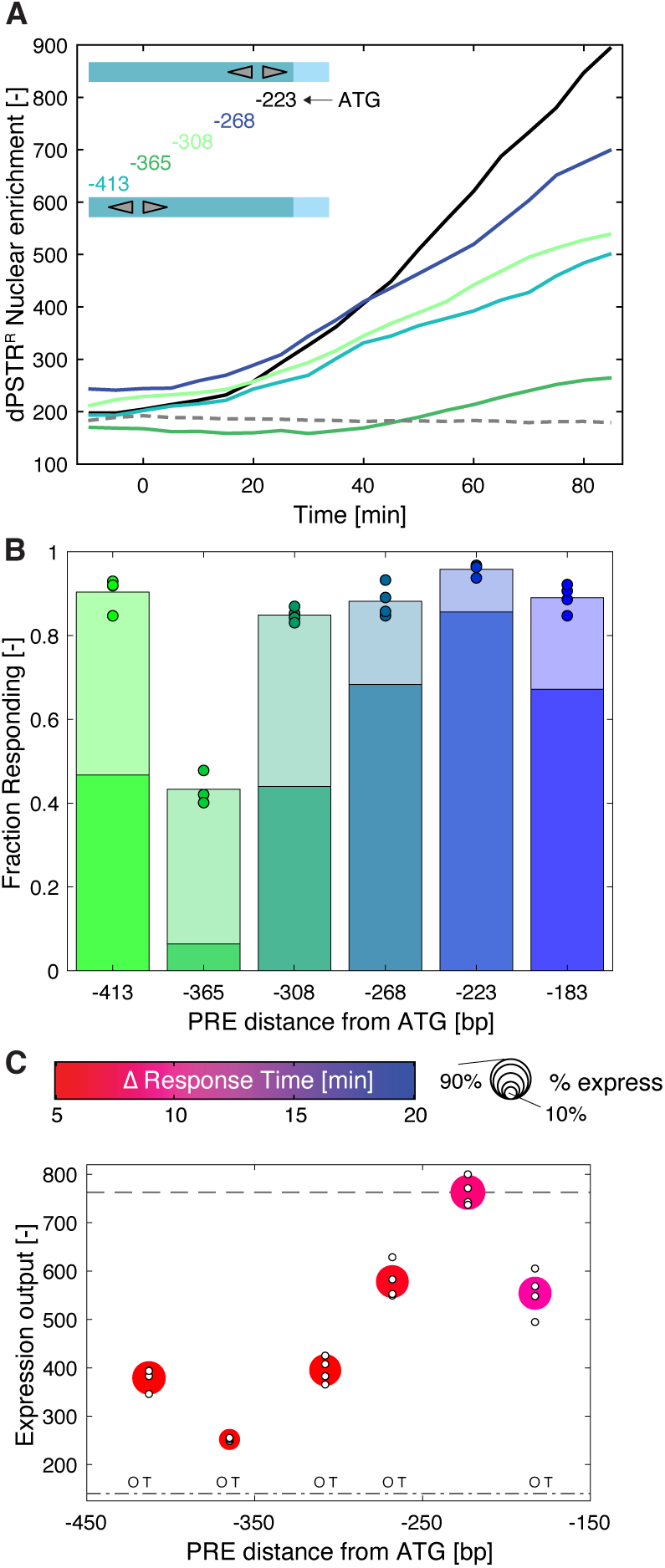
Distance of the PREs to the core promoter impacts expression output in the synthetic promoter. A. Dynamics of nuclear relocation for the dPSTR^R^ controlled by two PREs spaced by 3 bp in tail- to-tail orientation placed at different positions on the promoter (color lines represent the median of the population). The solid black line represents the median response of the PREs placed 223 bp downstream of the start codon (p*SYN*3TT), while the dashed line represents the control promoter without PRE. B. Fraction of responding cells for the various PRE sequences. The darker portion of the bar represents the fraction of highly expressing cells (expression output > 50% of reference expression output) and the light bar the fraction of low-expressing cells (> 20% expression output < 50%). The marker represents the fraction of responding cells for individual replicates. C. Summary graph displaying the expression output, the speed and the fraction of responding cells for PRE placed at various distances from the Start codon. The color of the marker indicates the difference in response time between the synthetic promoter and the reference p*AGA1*-dPSTR^Y^. The size of the marker represents the fraction of responding cells. The expression output of individual replicates is indicated by small white dots. The dashed line represents the expression output and the dashed dotted line the expression threshold calculated based on the p*SYN*3TT. The O and T indicate a significant difference between the mean of the replicates (t-test: p-val < 0.05) in the timing of induction (T) or in the expression output (O) relative to the p*SYN*3TT.

### Interplay between Ste12 and nucleosomes

The modulation of the distance between core promoter and PRE was also tested using a modified *AGA1* promoter where all the consensus binding sites were mutated, and a PRE-dimer was moved from -128 to -435bp relative to the Start codon (Figure 5A). In this context, however, not only the expression levels but also the dynamics of induction were strongly influenced by the location of the PRE-sites (Figure 5B, C and D). We believe that this behavior can be explained by the position of the nucleosomes on the promoter. Indeed, the promoter activation is rapid and strong when the PREs fall in a nucleosome-depleted region (NDR). When the sites are located within a region protected by a nucleosome, the fraction of responding cells and the level and speed of induction are all attenuated. The most significant impact is observed with the PRE-dimers located at -330bp, where Ste12 binding is conflicting with a nucleosome centered around -349bp(43). Interestingly, the fraction of expressing cells for this promoter is low (18%), but the few cells that express display a substantial nuclear enrichment of the dPSTR^R^ (Figure 5E). This stochastic activation suggests the presence of a dynamic interplay between the binding of the nucleosomes and the Ste12 dimer. When the nucleosome is bound, no transcription takes place. In some cells, the histones can be displaced by the binding of Ste12, which remains stably associated to the promoter to induce a sizable but delayed expression.

**Figure 5.**
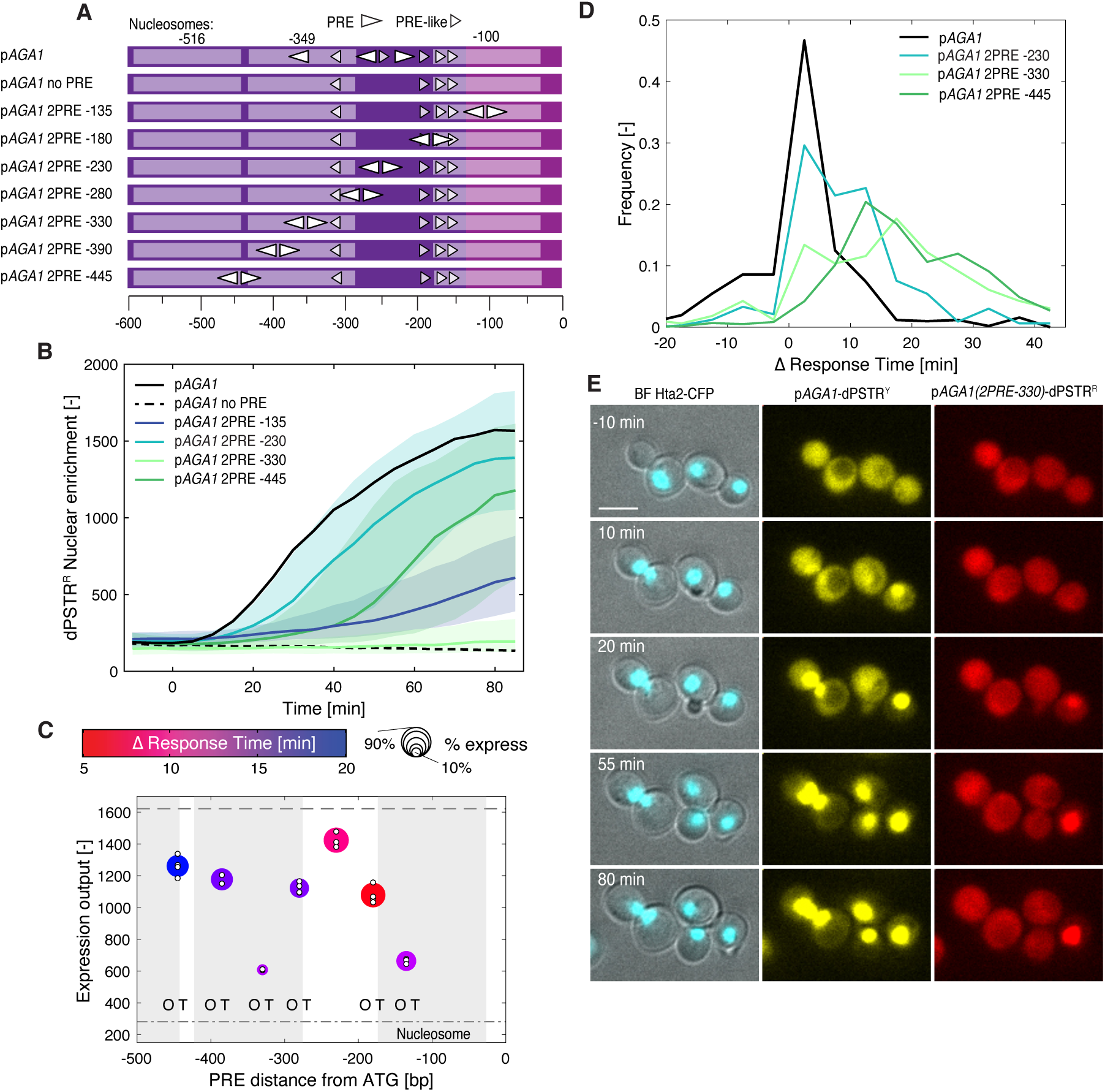
Varying the location of the PRE dimer in an endogenous promoter can influence both level and dynamics of induction. A. Scheme of the modified p*AGA1* promoters with consensus PRE sites indicated by elongated arrowheads and non-consensus PRE-like sites by small arrowheads. The regions protected by the nucleosomes positioned at -100, -349 and -516 are indicated by a lighter color. B. Dynamics of nuclear relocation for the dPSTR^R^ controlled for a selected set of modified p*AGA1* promoters containing two PRE spaced by 3 bp in tail-to-tail orientation placed at different positions along the promoter (colored lines median of the population and shaded area 25- to 75- percentile). The solid black line represents the median response of the reference endogenous AGA1 promoter, while the dashed line represents a mutated p*AGA1* where the three PRE and one PRE-like sites are mutated. C. Summary graph displaying the expression output, the speed and the fraction of responding cells for the complete set of the modified p*AGA1* promoters. The color of the marker indicates the difference in response time between the synthetic promoter and the reference p*AGA1*-dPSTR^Y^. The size of the marker represents the fraction of responding cells. The expression output of individual replicates is indicated by small white dots. The dashed line represents the expression output and the dashed dotted line the expression threshold calculated based on the p*AGA1*- dPSTR^R^. The gray areas represent the regions of the promoters covered by the three nucleosomes. The O and T indicate a significant difference between the mean of the replicates (t-test: p-val < 0.05) in the timing of induction (T) or in the expression output (O) relative to the p*AGA1* with 2 PRE positioned at -230. D. Histograms of the difference in response time between a few selected modified p*AGA1*-dPSTR^R^ and the reference p*AGA1*-dPSTR^Y^ in the cells that express both constructs. E. Thumbnail images of cells bearing the p*AGA1*-dPSTR^Y^ and the modified p*AGA1*-dPSTR^R^ with the PRE dimer positioned 330 base pairs before the Start codon. While all cells in the image relocate the dPSTR^Y^, only one cell displays a relocation of the dPSTR^R^. The scale bar represents 5 µm.

To test this model more directly, the promoter sequence was mutated to introduce a nucleosome disfavoring sequence (poly dA/dT, 20bp long) in the vicinity of the Ste12 binding sites (Figure 6A). The fraction of expressing cells increased from 20% to 50% when the dA/dT element was placed 6 bp away from the PRE sites (Figure 6B and C). An alternative option to increase the binding efficiency of Ste12 in the nucleosome bound region is to use multiple PRE sites. We used conformations with 3 PREs present in PRM1 and FUS1. The multiple sites and the high A/T content of the element facilitated the association of Ste12 to the promoter and resulted in an increased fraction of transcribing cells (Figure 6B and C). However, for all these constructs, the dynamics of induction were slow compared to the endogenous p*AGA1*. Interestingly, the addition of the dA/dT sequence slightly accelerated the induction of the reporter, suggesting that in a fraction of the population Ste12 can associate to the promoter under basal conditions (Figure 6D).

**Figure 6.**
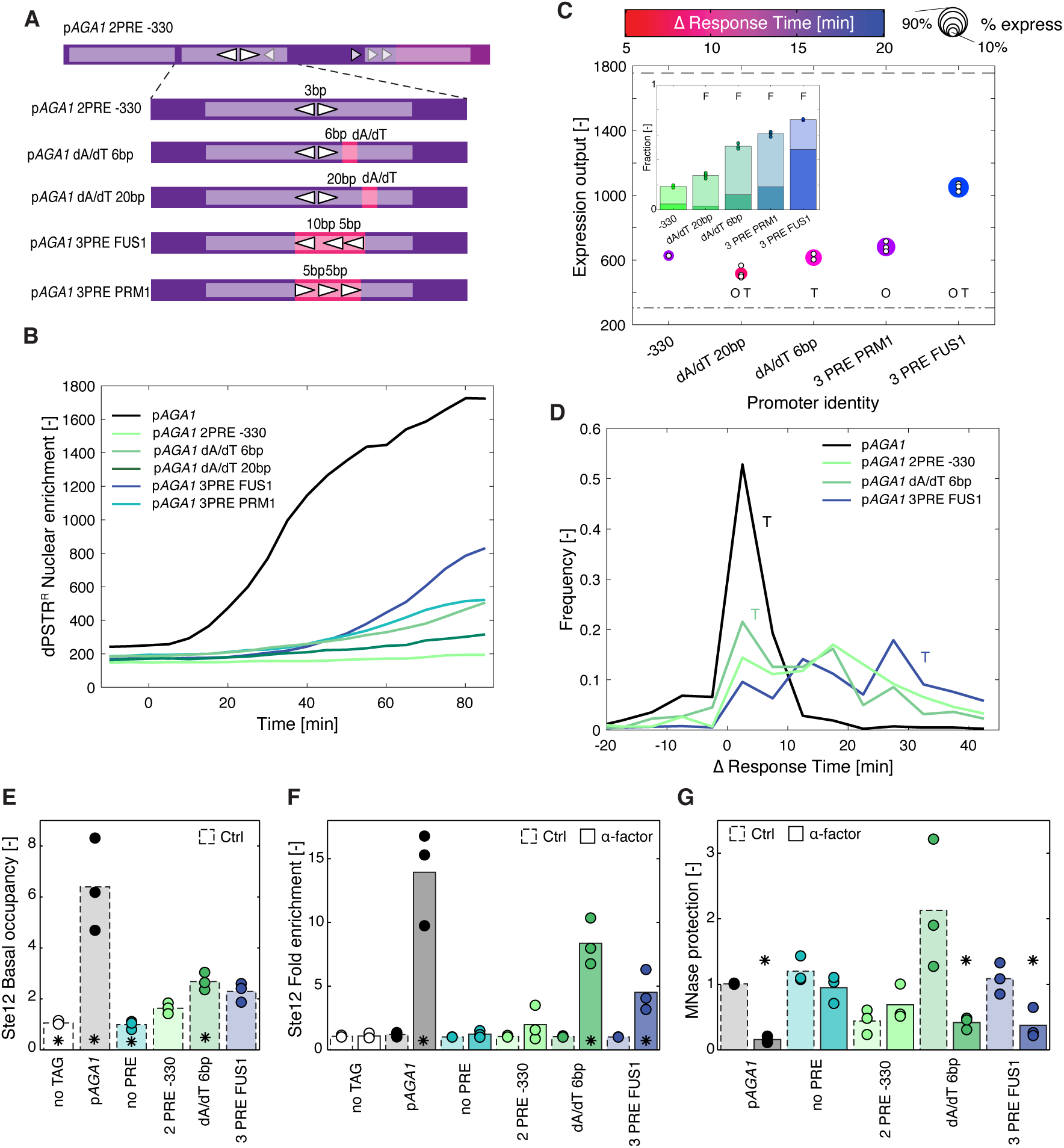
Favoring nucleosome eviction increases expression output. A. Scheme of the various promoters tested. Starting from the modified p*AGA1* with 2 PREs in tail-to-tail orientation at -330, poly-dA/dT segments of 20 bp were generated by mutations 6 or 20 bp away from the 2 PREs. Alternatively, the two PREs were replaced by three PRE sites extracted from the FUS1 or PRM1 promoters. B. Dynamics of nuclear relocation for the dPSTR^R^ controlled by modified p*AGA1* promoters. The four different modified promoters tested are shown in color. The solid black line represents the median response of the reference endogenous AGA1 promoter, while the dashed line represents the median of the population of cells with the 2 PREs in tail-to-tail orientation at -330. C. Summary graph displaying the expression output, the speed and the fraction of responding cells. The color of the marker indicates the difference in response time between the synthetic promoter and the reference p*AGA1*-dPSTR^Y^. The size of the marker represents the fraction of responding cells. The dashed line represents the expression output and the dashed dotted line the expression threshold calculated based on the p*AGA1*-dPSTR^R^. The O and T indicate a significant difference between the mean of the replicates (t-test: p-val < 0.05) in the timing of induction (T) or in the expression output (O) relative to the p*AGA1* with 2 PRE positioned at -330. The inset represents the fraction of responding cells for each promoter. The dark portion of the bar represents the fraction of highly expressing cells (Expression output > 50% EO of the reference promoter) and the light bar the fraction of low-expressing cells (20%< EO < 50%). The marker represents the fraction of responding cells for individual replicates. The F indicates that the fraction of responding cells is significantly different relative to the p*AGA1* with 2 PRE positioned at -330. D. Histograms of the response time difference measured between the modified p*AGA1*-dPSTR^R^ and the reference p*AGA1*-dPSTR^Y^. The solid black line corresponds to the endogenous AGA1 promoter, while the dashed line represents the 2 PREs in tail-to-tail orientation at -330. The addition of the poly-dA/dT 6bp away from the PRE dimer (light green) slightly accelerates the induction of the promoter. Placing the three PREs of FUS1 (blue) slows down the induction of the promoter. The T indicates that the histograms for the two mutated promoters are significantly different from the endogenous promoter using a Wilcoxon rank sum test. E. Basal association of Ste12 on various p*AGA1* promoters evaluated by Ch-IP in a region centered around -259bp. The values are normalized relative to the no TAG control. The star indicates a significant difference by a star (t-test: p-val < 0.05) relative to the promoter with 2 PRE at position -330. F. Fold induction in Ste12 binding measured by Ch-IP after 30 min treatment with pheromone. The star indicates a significant difference between the untreated control and the α-factor stimulated samples (t-test: p-val < 0.05). G. Eviction of nucleosomes monitored by MNase protection assays after 30min pheromone treatment in the same region as in E. Nucleosome occupancy is normalized relative to the untreated p*AGA1* endogenous promoter. A star indicates a significant difference between the untreated control and the α-factor stimulated samples (t-test: p-val < 0.05).

To probe the association of Ste12 on the DNA, Chromatin Immuno-Precipitation (Ch-IP) was performed on some of these promoter variants by amplifying a region centered around -259bp. The basal association of Ste12 is much higher in the endogenous *AGA1* sequence than in our promoter variants (Figure 6E). In addition, we see a small but significant enrichment of Ste12 on the promoter when the dA/dT stretch is added 6 bp away from the two PREs compare located at -330. Upon pheromone treatment Ste12 becomes more than 10-fold enriched on the endogenous promoter while we don’t detect a significant enrichment of the TF on the 2 PRE placed at -330 (Figure 6F). In contrast, promoters modified with the 3 PREs using the *FUS1* conformation or with the dA/dT stretch 6bp away from the PREs, the enrichment of Ste12 is significant. In parallel, the association of histones in the same region of these promoters was evaluated by MNase protection assays. On the WT p*AGA1*, we observe a strong eviction of the nucleosomes 30 minutes after the stimulation of the cells with α-factor (Figure 6G). No significant eviction can be observed on the promoter with the 2 PRE at -330 bp, while the eviction is recovered when the dA/dT stretch is added 6 bp away from the two PREs or if three PREs are present. Both the recruitment of Ste12 and the eviction of the nucleosomes are in line with the higher expression output measured with the dPSTR^R^. These biochemical data confirm the hypothesis that the access to the PRE sites at - 330 is limited for Ste12 under basal conditions which limits its ability to activate transcription from this site. However, the addition of a third PRE or the presence of a dA/dT stretch favors the access of Ste12 to the promoter by facilitating the eviction of the nucleosomes upon pheromone treatment.

The endogenous p*AGA1* promoter contains three consensus PREs and at least five PRE-like sites (Supplementary Figure 7A). The PRE/PRE-like sites spaced by 3 bp at position -220 alone seem sufficient to ensure a high induction level. Therefore, it is not clear why there are so many additional Ste12 binding sites present in this sequence. One of their roles might be to increase the local concentration of Ste12 in the vicinity of the locus. Interestingly, if the first PRE at -196 bp or its neighboring PRE-like at -185 bp are mutated, the level of induction of the promoter remains almost identical, while the dynamics of induction are delayed (Supplementary Figure 7B and C). Since these two PREs are 5bp apart in a head-to-tail conformation, it is likely that they allow the formation of a Ste12 dimer. One possible explanation for the delayed expression observed is that these two binding sites contribute to defining the nucleosome depleted region in the p*AGA1* promoter which favors the basal association of Ste12 to the central PRE of the promoter. Regardless, together, these results demonstrate that nucleosomes prevent the basal association of Ste12 outside of the NDR. Sites present in a nucleosome protected region of the promoter will display slow induction kinetics and a low fraction of inducing cells. Additional PRE sites on endogenous promoters may contribute to shaping the NDR and thereby allowing indirectly an early activation of transcription.

## Discussion

The coding sequence of a protein can provide insights into many of its properties. However, predicting the level and the timing of its expression based on the promoter sequence remains a challenge(44). Endogenous promoters display a large and complex palette of regulation combining binding sites for multiple TFs at various positions. Even in the simpler case of the mating pathway in budding yeast, where 200 genes are under the control of the single TF Ste12, the diversity in the organization of the PREs on these promoters is large. To elucidate the fundamental rules governing the induction of mating genes by Ste12, we designed synthetic promoters where the conformations of Ste12 binding sites could be tested systematically. To compare all these synthetic constructs, we placed them upstream of the core of the p*CYC1* promoter. Interestingly, a comparison between the p*AGA1* and the p*CYC1* core sequences indicate that this region also contributes to the regulation of the level and the speed of gene induction, which should be investigated further.

Based on these results we have identified various strategies that are at play in mating dependent promoters to regulate the properties of induction of a promoter. The level of induction can be controlled by two parameters: the binding affinity of the Ste12 sites and the distance from the core promoter. p*AGA1*, p*FAR1* and p*STE12* all possess 2 PRE spaced by 3bp in tail-to-tail orientation. The PRE dimers is placed at -200 bp from the start in p*AGA1* while it is located at -300 bp and - 400 bp for p*FAR1* and p*STE12* which could account for their lower inducibility(17). The strength of Ste12 association on a promoter can be tuned by point mutation in the PREs or by changing the binding conformation. For instance, the two PREs spaced by 13 bp (-250 bp from ATG) in p*SST2* results in lower induction compared to p*AGA1*. p*KAR4* and p*AGA1* both have a PRE dimer spaced by 3bp positioned ∼200 bp from the start codon with one of the PRE which contains two point-mutations(24). The stronger level of p*AGA1* induction suggests that the binding to the p*AGA1* PRE dimer is tighter. These comparisons clearly oversimplify the complexity of these promoters. All these endogenous sequences harbor numerous additional PRE or PRE-like sites and different core promoter sequences that together contribute to the final inducibility of the promoter. However, our findings allow the identification of which PRE and/or PRE-like on a promoter are in the proper configuration to allow the formation of a Ste12 dimer.

### Role of Kar4

Previous measurements have suggested that Kar4 interacts directly with the DNA and that late mating genes contain specific binding sites for Kar4(18). However, we favor a model where Kar4 is recruited to the promoters via its interaction with Ste12. Multiple evidences point in this direction. We have shown previously that Kar4 is recruited to the promoters of both early and late genes(17). The yeast epigenome project has identified a binding site for Kar4 on DNA which corresponds to the one of Ste12(43). Recent point mutants of Kar4 which display mating deficiencies have a decreased interaction with Ste12(45).

Our present data demonstrate that Kar4 has a dual effect on some of our synthetic promoters. First, deleting *KAR4* results in a delayed induction for promoters with lower affinity Ste12 binding sites. This suggests that Kar4 contributes to the stabilization of Ste12 on these weaker sites under basal conditions such that upon stimulation of the pathway, these promoters can be activated promptly. In parallel, we observe an increase in the level of induction from these same promoters containing PRE-like sites in *kar4Δ* cells. This surprising behavior could act via the activation domain of Ste12 since we don’t observe a similar increase in cells expressing the Ste12-EV chimeric TF in the absence of *KAR4*.

Absence of Kar4 slows down the induction of intermediate genes (*PRM1*, *FIG2*) and is required for the activation of late genes such as *FIG1* or *KAR3*(17). These late promoters bear PRE-like combined with PRE which are hidden by nucleosomes. In the absence of Kar4, the binding of Ste12 to the promoter is presumably too weak to be stabilized and displaced by the nucleosomes without resulting in productive transcription whenever it binds.

### Dynamics of promoter induction

We identified two strategies to achieve a slow induction of the promoter: either by restricting the access of Ste12 to the promoter by nucleosomes or by using an unconventional PRE dimer (23bp tail-to-tail or 15bp head-to-tail or head-to-head). In the two dozen mating-dependent promoters that we have analyzed, we identified only one with 15 bp spacing. Thus, the nucleosome protection may be the preferred option, because it allows tuning both the speed and level of induction. Indeed, the large spacing between two PRE sites can only achieve a slow and low level of gene expression. In contrast, when the PRE dimer is moved within the p*AGA1* promoter delayed but high induction can be generated. Positioning the PRE dimer under the nucleosome slows down the induction but also strictly reduces the fraction of responding cells. However, placing the binding sites closer to the edge of a nucleosome protected region seems to be sufficient to obtain a similar decrease in the speed of induction without affecting severely the expression level and, importantly, the fraction of responding cells. In the promoter of the late gene *FIG1*, a PRE dimer (5bp, tail-to-head) essential for gene expression is also placed at the boundary of the sequence protected by the nucleosome(17). This positioning of the PRE dimers delays the induction of *FIG1* until it is required for the fusion of the two mating cells without compromising its robust expression when it is needed. Positioning the PREs further into the nucleosome protected region could lead to a stochastic activation of the protein which could be detrimental for the mating outcome.

The timing of gene expression is crucial for many processes, from the rapid induction of stress response genes to the controlled induction of proteins during the cell cycle. Cell-fate decisions are characterized with a chronology of gene expression which requires the ability to tune both the level and the dynamics of gene expression independently of each other. Regulating the basal association of Ste12 on promoters via the positioning of nucleosomes offers the opportunity to control the chronology of gene expression during the mating process. With our improved understanding of the regulation of the mating gene expression, we are in a better position to perturb the timing of key proteins implicated in mating to verify the importance of this chronology on the mating process.

## Material and Methods

### Yeast strains and plasmids

The synthetic promoters were derived from a plasmid containing a slightly modified p*AGA1*- dPSTR^R^ plasmid(17) which contains a *ClaI* site at position 144 bp separating the regulatory region of *AGA1* (-1000 to -150) and the core region (-150 to 0). First the core promoter was replaced by the core promoter from *CYC1* (-180 to 0) cloned between *ApaI* and *ClaI*. Then the regulatory region (between *ClaI* and *AatII*) was replaced by fragments from a modified p*CYC1* promoter (138 bp) which destabilizes the association of nucleosomes(37). This fragment encodes various conformations of Ste12 binding sites. The distance to the Start mentioned in the text and figures corresponds to the distance to the end of the *CYC1* or *AGA1* promoters. The actual Start of the inducible dPSTR moiety is 53 bp downstream due to the presence of multiple restriction sites. The longer spacing between Start and PRE sites (Figure 4) were obtained by duplicating the *CYC1* regulatory region. The core promoter and the multiple UAS sequences were synthesized as double stranded DNA fragments (IDT) or as plasmids (GeneScript). The list of synthetic sequences is provided in Strain and Plasmids Supplementary File The plasmids obtained were verified by restriction digestion and sequencing and were transformed in the same reference strain from the W303 background with the histone Hta2 tagged with CFP (pGTH-CFP)(46) and containing the reference p*AGA1*-dPSTR^Y^ (17). The strains used in this study are listed in Strains Plasmids and Primers Supplementary File.

Gene deletions were performed using the pFA6a KAN cassette(47). Correct insertion of the deletion cassette was verified by PCR on genomic DNA. The human Estradiol receptor and the VP16 activation domain (EV)(39,40) were cloned in a pGT-NAT plasmid between *BamHI* and *NheI*. The Ste12-EV chimeric transcription factor was generated by transforming the EV sequence and NAT cassette with primers containing homology to the sequence flanking the activation domain of Ste12 (from 647 bp to STOP codon) in strains containing the selected synthetic promoters controlling the dPSTR^R^. The transformants were selected on SD-UHL + NAT plates. The correct insertion of the EV sequence in frame with the Ste12 ORF was controlled by sequencing PCR fragments obtained from genomic DNA. The synthetic transcription factor Z4- EV is placed under the control of the RPL30 promoter(48). The Venus is control by a modified p*GAL1* containing 6 Z4 binding sites. The pZ41BS and pZ42BS are based on the p*CYC1* promoter containing either one or two Z4 binding sites and driving the expression of the dPSTR^R^.

Typically, eight transformants were selected and screened visually to verify the expression level of the three fluorescent constructs (Hta2-CFP, p*AGA1*-dPSTR^Y^, p*SYN*-dPSTR^R^). Four transformants were then quantified in time-lapse experiments and used as biological replicates for the measurements of one promoter. Based on the consistency of the p*AGA1*-dPSTR^Y^ response and the behavior of the tested dPSTR^R^, we excluded some of the replicates. If two or more replicates were not providing consistent results, the experiment was repeated with different transformants. In each experiment, the behavior of 500 cells is typically quantified, but some replicates contain only 200 cells and others up to 1000 individual cells. In the dPSTR nuclear relocation graphs and in the response time histograms, we represent the behavior of one representative replicate. In the bar graphs displaying the fraction of responding cells or in the summary graphs, the mean behavior of all the selected replicates is plotted.

### Time-lapse microscopy

Yeast strains were grown overnight to saturation in SD-full medium (Complete CSM DCS0031, ForMedium) diluted in the morning in fresh SD-full and grown for at least 4 hours. The cultures were diluted to OD 0.04 and briefly sonicated and 200µl were loaded in the well of a 96-well plate (PS96B-G175, SwissCI) previously coated with Concanavalin A (L7647, Sigma-Aldrich). Cells settled in the well for 30 minutes before the start of the time-lapse.

Cells were imaged on an inverted wide-field epifluorescence microscope (Nikon, Ti2) enclosed in an incubation chamber set at 30° using a 40X Oil or a 40X AIR objective (Figure 2B to D Supplementary Figure 3E and F, Supplementary Figure 5E, F, Supplementary Figure 6C to F). The fluorescence excitation is provided by a Lumencor Spectra 3 light source (LED intensity 50%). A CFP YFP RFP dichroic filter (F68-003) and appropriate emission filters were used to detect the fluorescence emission using a sCMOS camera (Hamamatsu, Fusion BT), with 50ms, 50ms and 100ms exposure time respectively. Up to 8 wells were imaged in parallel and 5 fields of view per well were monitored. Two brightfield images (one slightly out of focus) and the three fluorescent images were recorded every 5 minutes for 100 minutes. Before the third time point, 100µl of a 3µM α-factor solution in SD-full was added to each well to reach a final concentration of 1µM in the well. The α-factor is a gift from the Peter lab at the ETHZ. To induce the Ste12-EV construct, 100µl of a 3µM β-estradiol (Sigma-Aldrich, E2758-250MG) solution in SD-full was used to reach a final concentration of 1µM in the well.

### Data analysis

Time-lapse images were segmented and single cell data extracted using the YestQuant platform(49). The CFP image allowed to determine the position of the nuclei of each cell. Using this first object, the two brightfield images were used to identify the border of each cell and define the cytoplasm object by removing the pixels belonging to the nucleus.

The quantification of the single cell traces was performed with Matlab (The Mathworks, R2023b). Only cells tracked from the beginning to the end of the timelapse were kept for the analysis. The nuclear enrichment is calculated as the difference between the average fluorescence of the nucleus object and the cytoplasm object. The basal nuclear enrichment is measured as the mean of the three first time points. The Expression Output (EO) corresponds to the difference between the maximum of a single cell traces and the basal level (Supplementary Figure 2D).

To determine if a cell is deemed transcribing or not, two criteria are used. First, the last 5 points of the trace minus the basal level has to be significantly higher than zero (sign-test. P-value 0.05). Second, the Expression output of the trace (maximum of the trace – basal level) has to exceed a threshold. To define the threshold, we select one strain as the reference and calculate the threshold as 20% of the mean Expression ouput from all cells. The strain bearing the p*AGA1*-dPSTR^R^ (Figures 5 and 6 and Supplementary Figures 2, 3 and 7) or the p*SYN*3TT-dPSTR^R^ (Figure 1, 3, 4 and Supplementary Figure 4) were used as references. Transcribing cells are further differentiated in strong and weak expression if their output exceed 50% of the Expression output of the reference strain and as weakly expressing if the expression output falls between 20% to 50% of the reference (Supplementary Figure 2D, E and F).

The response time is defined as the first time point after the trace overcomes 20% of its own expression output (Supplementary Figure 2D). The difference in response time between the tested dPSTR^R^ and the internal reference construct p*AGA1*-dPSTR^Y^ is only calculated for cells considered as transcribing both reporters (Supplementary Figure 2G and H).

### ChIP assays

Yeast cultures were grown to early log phase (A660 0.4–0.6), then samples (50ml) were subjected to 1µM α-factor for 30 minutes. For crosslinking, yeast cells were treated with 1% formaldehyde for 20 minutes at room temperature. Glycine was added to a final concentration of 330 mM for 15 minutes. Cells were collected, washed four times with cold TBS (20 mM Tris-HCl, pH 7.5, 150 mM NaCl), and kept at −20 °C for further processing. Cell pellets were resuspended in 0.3 ml cold lysis buffer (50 mM HEPES-KOH, pH 7.5, 140 mM NaCl, 1 mM EDTA, 0.1% sodium deoxycholate, 1% Triton-X 100, 1 mM PMSF, 2 mM benzamidine, 2 µg/mL leupeptin, 2 µg/mL pepstatin, 2 µg/mL aprotinin). An equal volume of glass beads was added, and cells were disrupted by vortexing (with Vortex Genie) for 13 minutes at 4°C. Glass beads were discarded and the crosslinked chromatin was sonicated with water bath sonicator (Bioruptor) to yield an average DNA fragment size of 350 bp (range, 100–850 bp). Finally, the samples were clarified by centrifugation at 16,000g for 5 minutes at 4°C. Supernatants were incubated with 50µL anti-HA 12CA5 monoclonal antibodies pre-coupled to pan mouse IgG DynabeadsTM (Invitrogen, 11042). After 120 minutes at 4°C on a rotator, beads were washed twice in 1mL lysis buffer, twice in 1mL lysis buffer with 500 mM NaCl, twice in 1mL washing buffer (10 mM Tris-HCl pH 8.0, 0.25 M LiCl, 1 mM EDTA, 0.5% N-P40, 0.5% sodium deoxycholate) and once in 1mL TE (10 mM Tris-HCl pH 8.0, 1 mM EDTA). Immunoprecipitated material was eluted twice from the beads by heating for 10 minutes at 65 °C in 50 µl elution buffer (25 mM Tris-HCl pH 7.5, 1 mM EDTA, 0.5% SDS). To reverse crosslinking, samples were adjusted to 0.3 ml with elution buffer and incubated overnight at 65°C. Proteins were digested by adding 0.5mg/ml Proteinase K (Novagen, 71049) for 1.5 hours at 37°C. DNA was extracted with phenol-chloroform-isoamyl alcohol (25:24:1) and chloroform. It was finally precipitated with 48% (v/v) of isopropanol and 90 mM NaCl for 2 hours at −20 °C in the presence of 20 µg glycogen, and resuspended in 30 µL of TE buffer. Quantitative PCR analysis on p*AGA1*-dPSTR-R used the following primers with locations indicated by the distance from the respective ATG initiation codon: *AGA1* promoter (- 310/-207); and *TEL* (telomeric region on the right arm of chromosome VI). Experiments were done on three independent chromatin preparations and quantitative PCR analysis was done in real time using an Applied Biosystems Via7. Immunoprecipitation efficiency was calculated in triplicate by normalizing the amount of PCR product in the immunoprecipitated sample by that in *TEL* sequence control. The binding data are presented as fold induction with respect to the non-treated condition, for basal binding of Ste12 the data are referenced to the untagged strain (no tag) which was set to 1.

### MNase nucleosome mapping

Yeast spheroplast preparation and micrococcal nuclease digestions were performed as described previously with modifications (50,51). Ste12-6xHA tagged strain was grown to early log phase (A660 0.4–0.6) and samples of 500 ml of culture were exposed to 1 µM α-factor for 30 minutes. The cells were cross-linked with 1% formaldehyde for 15 minutes at 30°C and the reaction was stopped with 125 mM glycine for minutes. Cells were washed and resupended in 1M sorbitol TE buffer before cell wall digestion with 100 T zymoliase (USB). Cells were then lysed and immediately digested with 60–240 mU/µl of micrococcal nuclease (Worthington Biochemical Corporation, Lakewood; NJ., USA). DNA was subjected to electrophoresis in a 1.5% (w/v) agarose gel and the band corresponding to the mononucleosome was cut and purified using a QIAquick gel extraction kit (Qiagen). DNA was used in a real-time PCR with specific tiled oligonucleotides covering the *AGA1* promoter of the dPSTR^R^ (the endogenous AGA1 gene has been mutated). PCR quantification was referred to an internal loading control (telomeric region in chromosome 6) and nucleosome occupancy was normalized to 1 at the (-1) nucleosome region of the untreated condition. The degree of nucleosome eviction at the indicated region was set to 1 in the wild type strain in control conditions and used as a reference.

## Data Availability

Analyzed single cell measurements are available on Zenodo (DOI: 10.5281/zenodo.14438716). The raw images of this study are available from the corresponding author upon reasonable request.

## Code Availability

The image analysis platform has been published previously^8^. A more recent version of the code can be obtained from the corresponding author. The analysis scripts are provided on the Zenodo repository along with the data.

## Acknowledgment

We thank members of the Pelet lab for critical comments on the project. We thank Michael Taschner and Stephan Gruber for helpful discussions, Marta Schmitt, Stella Parzanese Yaima Matas and Mònica Romo for technical help

## Author Contributions

SPi and SPe designed the experiments and wrote the manuscript. SPi performed the time-lapse measurements and SPe analyzed the data. CS and MNR performed the biochemistry experiments and FP and MNR analyzed the results. SPi, VV and YD generated the strains and plasmids used in the study.

## Supplementary Information Captions

- Supplementary Figures 1 to 7.

- Data File S1: Listing every yeast strains, plasmids and primers used in this study.

