## Supplementary Figures 1 to 7 for "Basal association of a transcription factor favors early gene expression"

### Supplementary Figure Legends

#### Supplementary Figure 1. Scheme of the mating pathway

Yeast cells detect pheromone via a G-protein coupled receptor. The G-protein disassemble and recruits the scaffold Ste5 at the plasma membrane. Then, Ste20 activates the MAP3K Ste11, which phosphorylates the MAP2K Ste7. Ste7 phosphorylates both MAPK Fus3 and Kss1. Fus3 phosphorylates Far1 to arrest the cell cycle in G1. Both Kss1 and Fus3 contribute to the transcriptional response by inhibiting the repression exerted on the TF Ste12 by Dig1 and Dig2.

#### Supplementary Figure 2. Quantification of the expression response.

A. Dynamics of nuclear enrichment of the dPSTR$^R$ under the control of the p*AGA1* (dark blue), the p*FIG1* (magenta) or the synthetic promoter with two PREs in tail-to-tail orientation with 3 bp spacing (p$SYN_{3TT}$ - cyan). The solid line represents the median of the population, while the shaded area represents the 25- to 75-percentiles of the population.

B. Dynamics of nuclear enrichment of the dPSTR$^Y$ under the control of the endogenous p*AGA1* promoter which is present in parallel to the test dPSTR$^R$ reporter for the three strains presented in panel A. The response of the p*AGA1*-dPSTR$^Y$ serves as a control for the robustness of pheromone induction for all experiments. If the p*AGA1*-dPSTR$^Y$ is not induced properly, the experiment will be rejected.

C. Correlation of the normalized expression level at 0, 20, 40, 60 min after the stimulus between the p*AGA1*-dPSTR$^Y$ (y-axis) and p*AGA1* (left), p$SYN_{3TT}$ (middle) and the p*FIG1* (right)-dPSTR$^R$ (x-axis). The Spearman correlation coefficient for each distribution is indicated in the lower right corner.

D. Description of the metrics measured from a single cell trace of nuclear enrichment. The mean of the nuclear enrichment of the first 3 time points is used to quantify the basal level of expression of the trace. The difference between the maximum of the trace and the basal level represents the expression output (EO). When the trace overcomes the threshold set by the 20% of this EO added to the basal level, the response time (RT) is defined.

E. To characterize individual single cell traces as not responding, weakly or strongly responding, the mean EO of all the cells of a reference strain is used. In the present case, the reference strain is the p*AGA1*-dPSTR$^R$ construct which is used as a reference. Two criteria are used to define expressing cells. First, the last 5 points of the trace have to be significantly higher than the basal level (sign-test, blue, red, yellow traces). Second, the Expression Ouput of the trace has to overcome the expression threshold set at 20% of the reference trace EO (blue, red, green). The yellow and green traces which satisfy only one condition are thus considered as not expressing.

In addition, if the expression output of a single cell exceeds 50% of the reference EO, it is considered as a strongly expressing cell (blue), while if it falls between the 20% to 50% it is defined as weakly expressing (red).

F. Fraction of responding cells for the dPSTR$^R$ (left) and the dPSTR$^Y$ (right). The dark bar represents the mean fraction of strongly responding cells and the light bar the weakly expressing ones. The round markers represent the total fraction of responding cells measured in the 2 to 4 biological replicates used to build the graph.

G. Scheme describing the calculation of the difference in response time between the p*AGA1*-dPSTR$^Y$ (top panels) and the test p*SYN*$_{3TT}$-dPSTR$^R$. In the left panels, three traces are shown with various delays in p*AGA1*-dPSTR$^Y$ induction and matching dynamics in the dPSTR$^R$ resulting in the calculation of a small difference in response time (ΔRT). On the right panels, two traces in the p*AGA1*-dPSTR$^Y$ display a fast response while the dPSTR$^R$ responses rise much later, resulting in a large ΔRT.

H. Histogram of the difference in response time (ΔRT) calculated for all the cells expressing both the reference p*AGA1*-dPSTR$^Y$ and the test dPSTR$^R$ controlled by p*AGA1* (blue), p*SYN*$_{3TT}$ (cyan) and p*FIG1* (magenta).

**Supplementary Figure 3. Development of a synthetic mating-responsive promoter.**

A. Scheme of the various promoter configurations tested starting from the p*AGA1* endogenous reporter and exchanging the core promoter and testing regulatory regions with a UAS containing various configurations of PREs.

B. Dynamics of nuclear enrichment for the dPSTR$^R$ under the control of various synthetic promoters. The colored solid lines represent the median of the population and the shaded area, the 25- and 75-percentile of the population. The solid black line is the reference induction from the endogenous promoter p*AGA1* and the dashed line is the control promoter without PRE sites.

C. Fraction of strongly (dark bar) and weakly (light bar) responding cells relative to the p*AGA1*-dPSTR$^R$. The total fraction of responding cells from individual replicates is displayed by the markers.

D. Histogram of the difference in response time between the tested promoter and the internal reference provided by the p*AGA1*-dPSTR$^Y$.

E. Dynamics of nuclear enrichment for the p*AGA1*-dPSTR$^Y$ (left) and p*SYN*-dPSTR$^R$ (right) variants in WT (solid line) and *ste12Δ* (dashed lines) strains.

F. Fraction of responding cells in the WT and *ste12Δ* strains for the p*AGA1*-dPSTR$^Y$ (left) and p*SYN*-dPSTR$^R$ (right) variants.

**Supplementary Figure 4. Influence of PRE orientation and spacing on the output of the synthetic promoter.**

A, B and C. Time course of the nuclear enrichment of the dPSTR$^R$ for various distances of PRE placed in tail-to-head conformation towards the core (A), in tail-to-head conformation away from the core (B) and in head-to-head conformation (C). Three spacings are plotted in color. The solid lines represent the median and the shaded area the 25- to 75- percentile of the population. Gray lines represent the median of non-functional PRE conformations. The solid black line represents the median of the p$SYN_{3TT}$ reference promoter. The black dashed line is the median of the control synthetic promoter without PRE.

D, E and F. Summary graph displaying the expression output, the speed and the fraction of responding cells for various spacings of the two PREs placed in tail-to-head conformation towards the core (D), in tail-to-head conformation away from the core (E) and in head-to-head conformation (F). The color of the marker indicates the difference in response time between the synthetic promoter and the reference p*AGA1*-dPSTR$^{Y}$. The size of the marker represents the fraction of responding cells. The dashed line represents the expression output and the dashed dotted line the expression threshold calculated based on the p$SYN_{3TT}$. The O and T indicate a significant difference between the mean of the replicates (t-test: p-val < 0.05) in the timing of induction (T) or in the expression output (O) relative to the p$SYN_{3TT}$.

**Supplementary Figure 5. Influence of binding site number for the β-estradiol-dependent induction by Z4-EV or Ste12-EV.**

A. Dynamics of nuclear enrichment of the dPSTR$^{R}$ under the control of synthetic promoters with one (blue) or two (green) Z4 binding sites (McIsaac *NAR* 2013) using the synthetic transcription factor Z4-EV upon stimulation with 1$\mu$m β-estradiol at time 0.

B. Increase in cellular fluorescence as function of time for the reference promoter containing 6 Z4 binding sites and driving the expression of a Venus fluorescent protein which serves as an induction control for the experiment plotted in panel A.

C. Dynamics of nuclear enrichment of the dPSTR$^{R}$ by the Ste12-EV upon stimulus by β-estradiol at time 0 under the control of different promoters containing zero (dark green), 1 PRE (light green) or two PREs in tail-to-tail orientation (blue). No detectable nuclear enrichment is observed for the 0 or 1 PRE controls, as well as, the 2 PRE spaced by 40 bp (light blue). If the 2 PREs are spaced by 3 bp (p$SYN_{3TT}$), the induction is strong (dark blue).

D. Dynamics of nuclear enrichment of the control p*AGA1*-dPSTR$^{Y}$ by the Ste12-EV in the strains containing the synthetic promoters displayed in panel C.

E. Comparison of the inducibility of p*AGA1*-dPSTR$^{Y}$ with the Ste12 WT (solid borders) or the Ste12-EV (dashed borders). The Expression Outputs for the p*AGA1* reporters induced by the Ste12 WT or the Ste12-EV were normalized relative to the Expression Output of the reference p$SYN_{3TT}$ sample. The bar represents the mean response of the replicates shown by the circles. A significant difference between the normalized EO Ste12-WT and Ste12-EV is indicated by a star (t-test: p-val < 0.05)

F. Fraction of responding cells for the p*SYN*-dPSTR$^{R}$ variants with the Ste12 WT (solid borders) or the Ste12-EV (dashed borders). The bar represents the mean response of the replicates shown by the circles. A significant difference between the fraction of responding cells between Ste12-WT and Ste12-EV is indicated by a star (t-test: p-val < 0.05).

**Supplementary Figure 6. Effect of the deletion of KAR4 on mating gene induction.**

A. Dynamics of nuclear enrichment of the p*AGA1*-dSPTR-Y in WT (solid lines) and *kar4Δ* cells (dashed lines).

B. Histograms of the difference in response time between the tested promoter and the internal p*AGA1*-dPSTR$^{Y}$ reference for WT (solid lines) and *kar4Δ* cells (dashed lines) for two different non-consensus PRE sequences associated to one consensus PRE.

C. Summary graph displaying the expression output, the speed and the fraction of responding cells for promoters with various PRE conformations in WT and *kar4Δ* cells. The color of the marker indicates the difference in response time between the synthetic promoter and the reference p*AGA1*-dPSTR$^{Y}$. The size of the marker represents the fraction of responding cells. The expression output of individual replicates is indicated by small white dots. The dashed line represents the expression output and the dashed dotted line the expression threshold calculated based on the p$SYN_{3TT}$ in WT cells. The O and T indicate a significant difference between the mean of the replicates (t-test: p-val < 0.05) in the timing of induction (T) or in the expression output (O) between the WT and *kar4Δ* strains for the same promoter.

D. Dynamics of nuclear enrichment of the p*SYN*-dPSTR$^{R}$ variants (right panel) and p*AGA1*-dSPTR-Y (left panel) in WT (solid lines) and *kar4Δ* cells (dashed lines).

E. Dynamics of nuclear enrichment of the p*SYN*-dPSTR$^{R}$ with two PRE spaced by 3 bp in tail to tail orientation with one mutated PRE (TcAAAC) in WT (solid lines) and *kar4Δ* cells (dashed line) with the chimeric Ste12-EV promoter and stimulated with $\beta$-estradiol at time 0.

F. Expression output of the strains measured in panel C. The O indicates that the pSYN$_{3TT}$ expresses significantly stronger than the two strain with the mutated PRE, which both express to the same level.

**Supplementary Figure 7. Mutation of the PRE1 and its associated PRE-like in p*AGA1* slows down response time.**

A. Scheme of the p*AGA1* endogenous promoter, which contains three consensus PRE sites and at least five non-consensus ones. PRE2 together with a non-consensus PRE spaced by 3 bp in tail-to-tail orientation are essential for the inducibility of the promoter. PRE1 (closest to the core) is spaced by 5 bp from a non-consensus site in tail to head conformation. PRE1 or its associated PRE-like have been mutated.

B. Dynamics of nuclear enrichment of the dPSTR$^{R}$ under the control of the endogenous (dark blue) or the mutated (light blue or magenta) *AGA1* promoter. The solid line represents the median of the population and the shaded area the 25- 75- percentile of the population.

C. Histogram of the difference in response time between the tested promoters and the internal p*AGA1*-dPSTR$^{Y}$ reference. The T indicates that the histograms for the two mutated promoters are significantly different from the endogenous promoter using a Wilcoxon rank sum test.

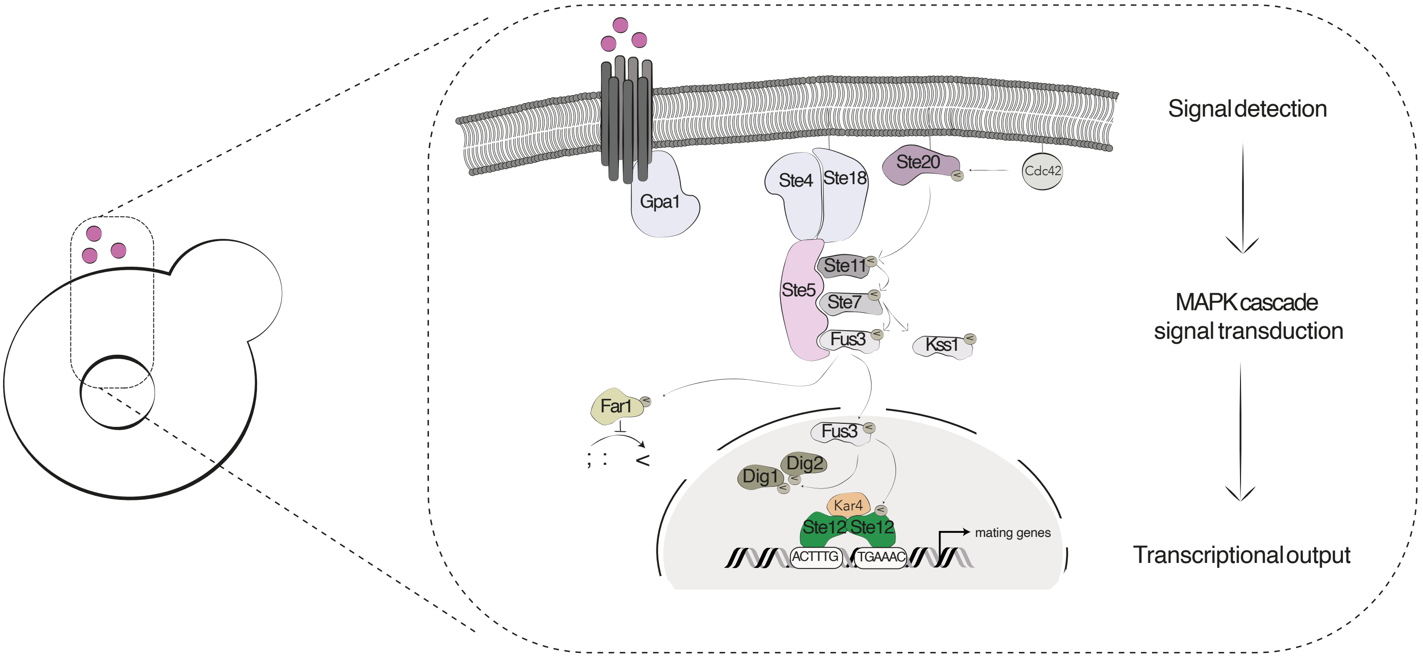


Supplementary Figure 1

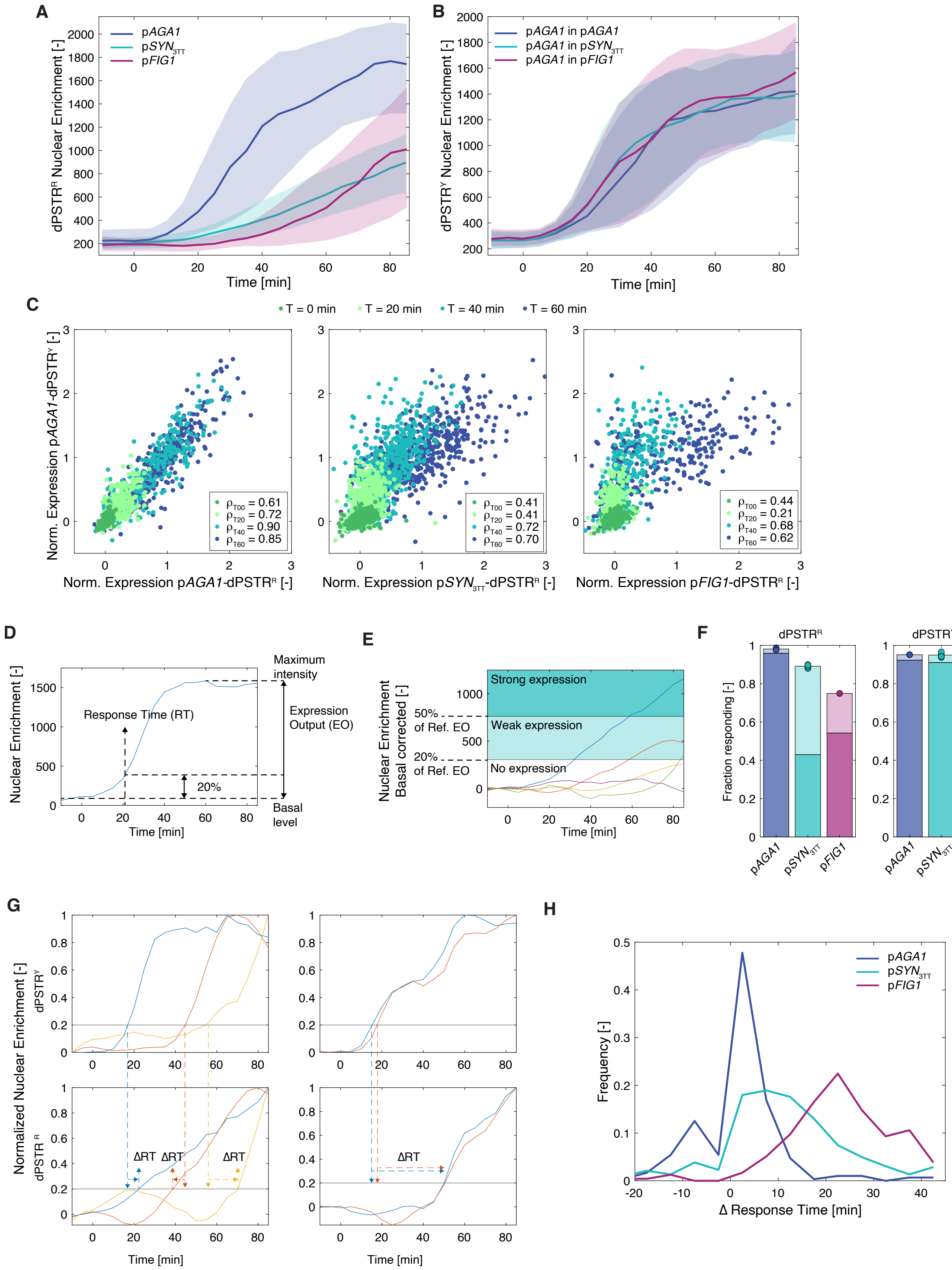


Supplementary Figure 2

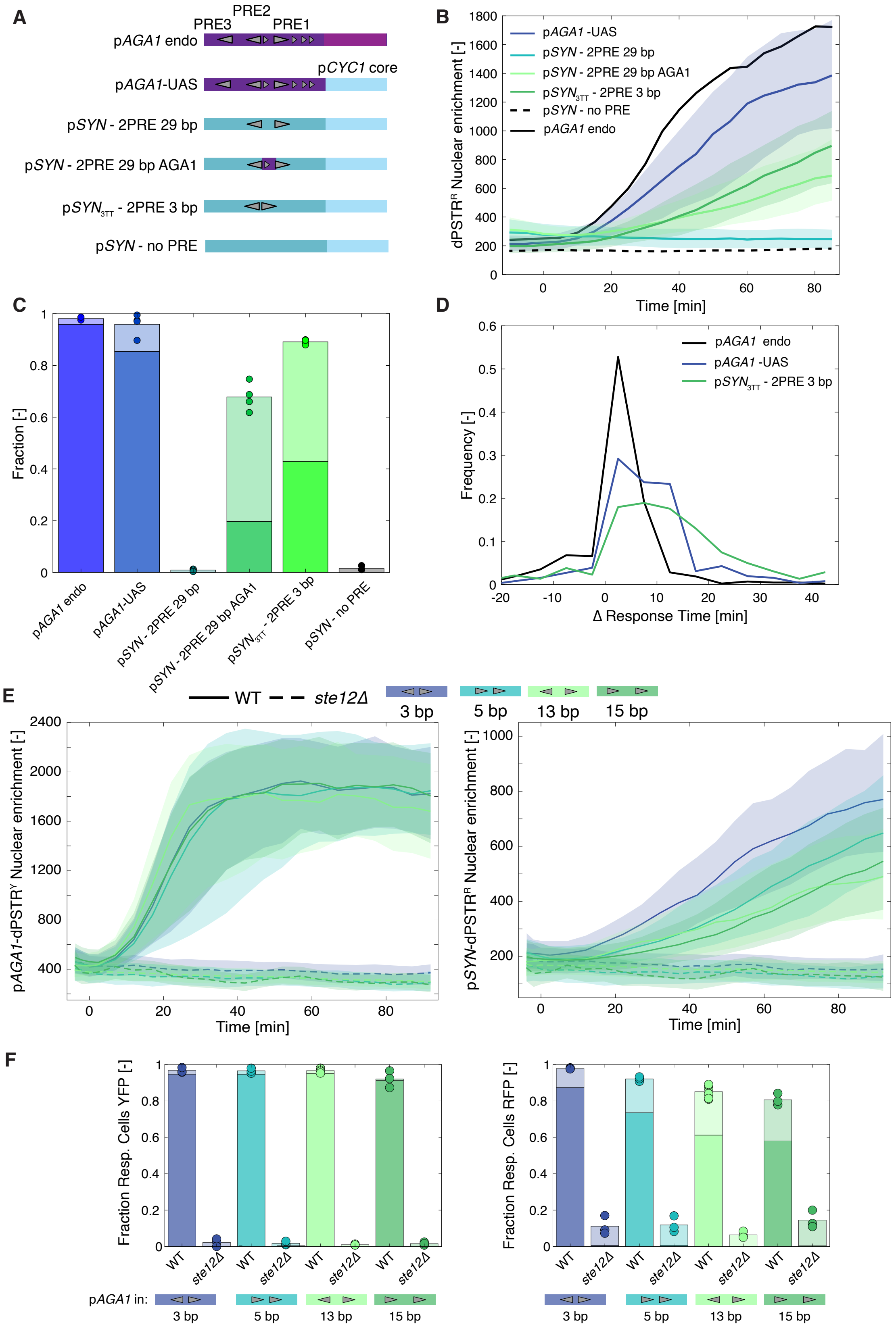


Supplementary Figure 3

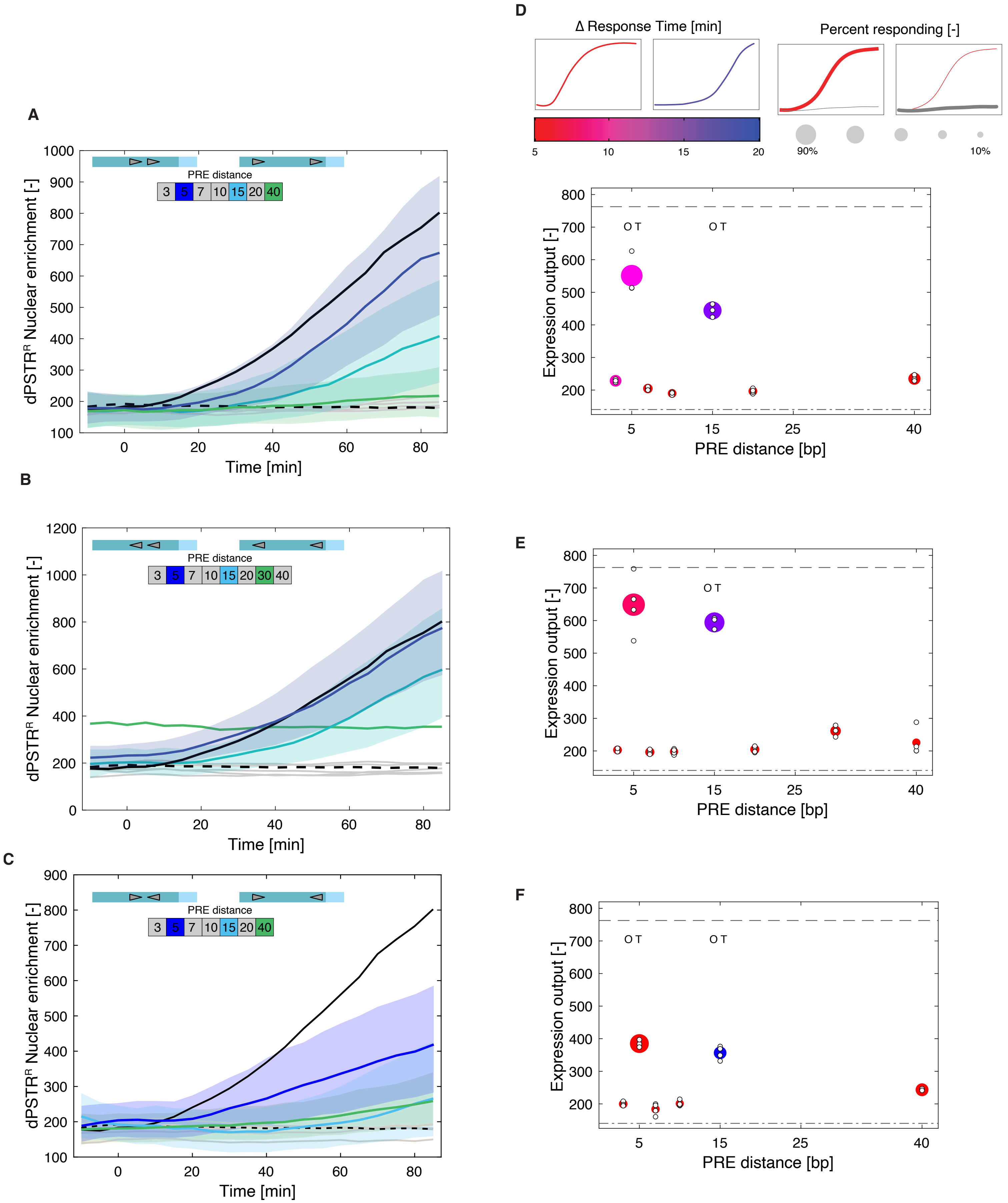


Supplementary Figure 4

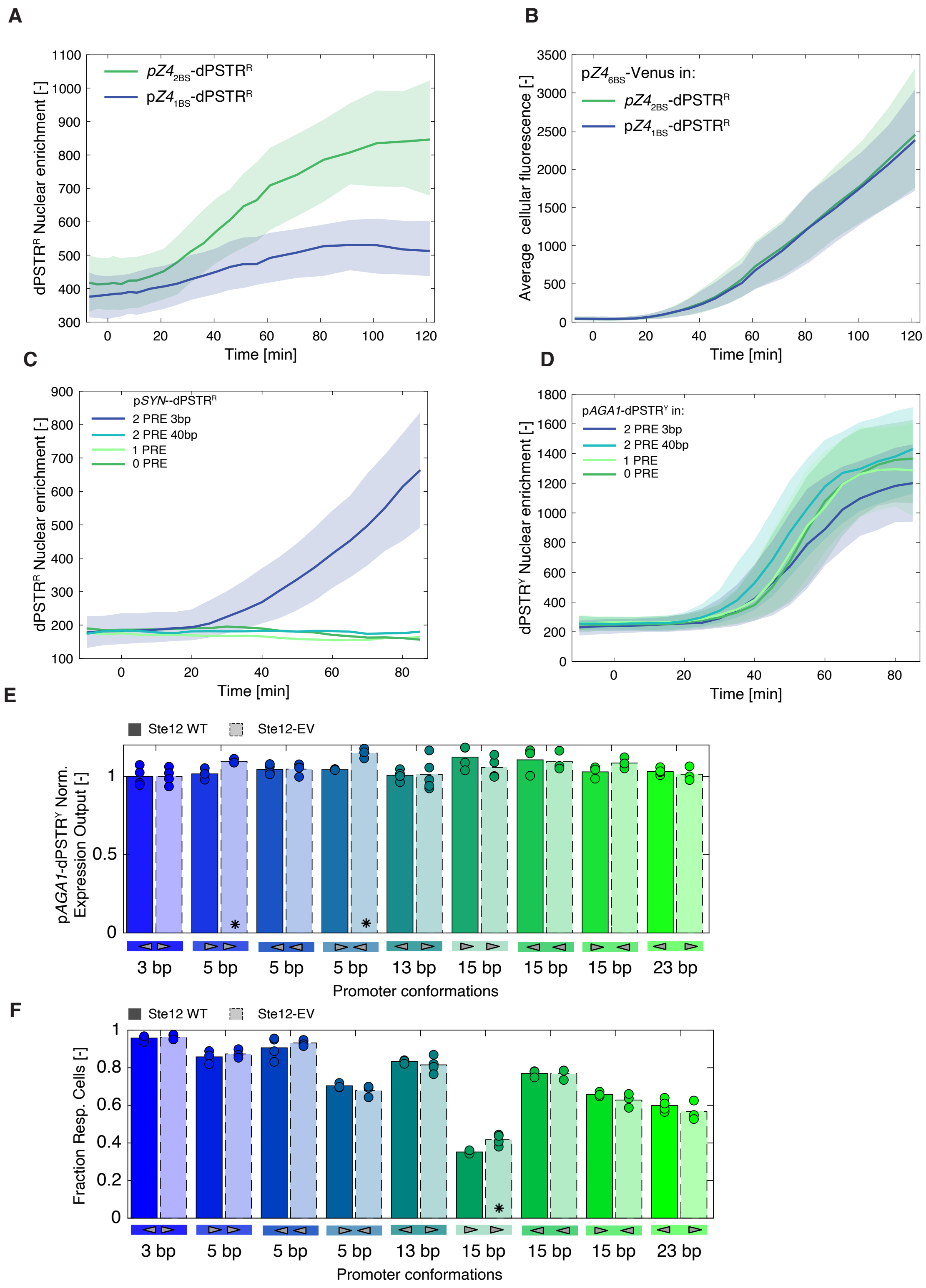


Supplementary Figure 5

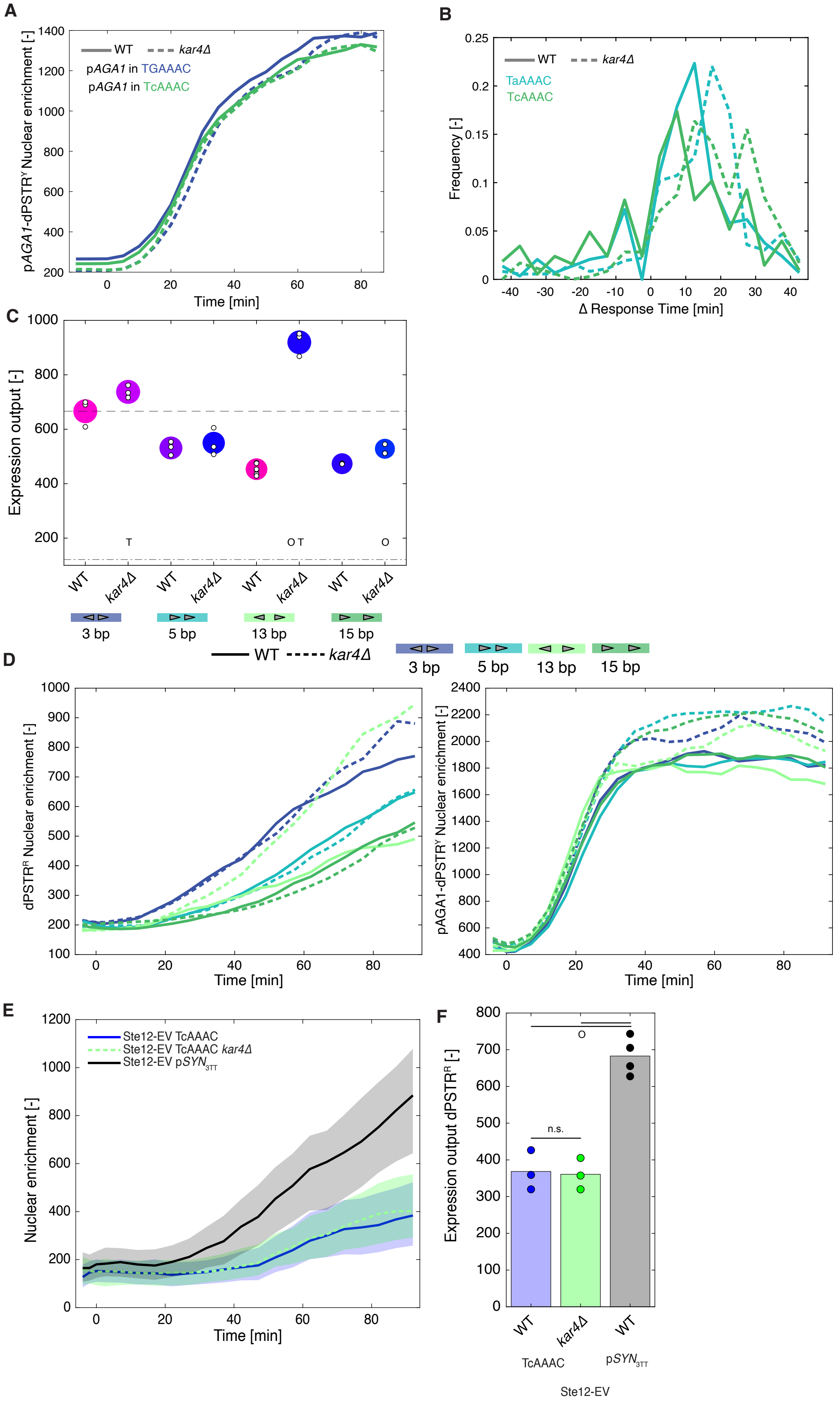


Supplementary Figure 6

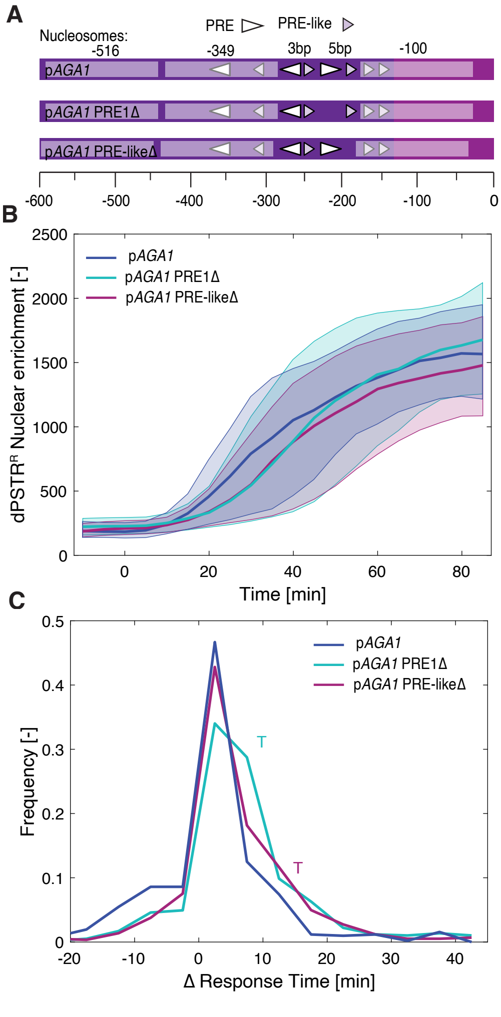


Supplementary Figure 7
